# Accurate prediction of *cis*-regulatory modules reveals a prevalent regulatory genome of humans

**DOI:** 10.1101/2020.05.15.098988

**Authors:** Pengyu Ni, Zhengchang Su

## Abstract

Annotating all *cis*-regulatory modules (CRMs) and transcription factor (TF) binding sites(TFBSs) in genomes remains challenging. We tackled the task by integrating putative TFBSs motifs found in available 6,092 datasets covering 77.47% of the human genome. This approach enabled us to partition the covered genome regions into a CRM candidate (CRMC) set (56.84%) and a non-CRMC set (43.16%). Intriguingly, like known enhancers, the predicted 1,404,973 CRMCs are under strong evolutionary constraints, suggesting that they might be *cis*-regulator. In contrast, the non-CRMCs are largely selectively neutral, suggesting that they might not be cis-regulatory. Our method substantially outperforms three state-of-the-art methods (GeneHancers, EnhancerAtlas and ENCODE phase 3) for recalling VISTA enhancers and ClinVar variants, as well as by measurements of evolutionary constraints. We estimated that the human genome might encode about 1.46 million CRMs and 67 million TFBSs, comprising about 55% and 22% of the genome, respectively; for both of which, we predicted 80%. Therefore, the *cis*-regulatory genome appears to be more prevalent than originally thought.

## Introduction

*cis*-regulatory sequences, also known as *cis*-regulatory modules (CRMs) (i.e., promoters, enhancers, silencers and insulators), are made of clusters of short DNA sequences that are recognized and bound by specific transcription factors (TFs)[1]. CRMs display different functional states in different cell types in multicellular eukaryotes during development and physiological homeostasis, and are responsible for specific transcriptomes of cell types[2]. A growing body of evidence indicates that CRMs are at least as important as coding sequences (CDSs) to account for inter-species divergence[3, 4] and intra-species diversity[5], in complex traits. Recent genome-wide association studies (GWAS) found that most complex trait-associated single nucleotide polymorphisms (SNPs) do not reside in CDSs, but rather lie in non-coding sequences (NCSs)[6, 7], and often overlap or are in linkage disequilibrium (LD) with TF binding sites (TFBSs) in CRMs[8]. It has been shown that complex trait-associated variants systematically disrupt TFBSs of TFs related to the traits [8], and that variation in TFBSs affects DNA binding, chromatin modification, transcription[9-11], and susceptibility to complex diseases[12, 13] including cancer[14-17]. In principle, variation in a CRM may result in changes in the affinity and interactions between TFs and their binding sites, leading to alterations in histone modifications and target gene expressions in relevant cells[18, 19]. These alterations in molecular phenotypes can lead to changes in cellular and organ-related phenotypes among individuals of a species[20, 21]. However, it has been difficult to link non-coding variants to complex traits[18, 22], largely because of our lack of a good understanding of all CRMs, their constituent TFBSs and target genes in genomes[23].

Fortunately, the recent development of ChIP-seq techniques for locating histone marks[24] and TF bindings in genomes in specific cell/tissue types[25] has led to the generation of enormous amount of data by large consortia such as ENCODE[26], Roadmap Epigenomics[27] and Genotype-Tissue Expression (GTEx)[28], as well as individual labs worldwide[29]. These increasing amounts of ChIP-seq data for relevant histone marks and various TFs in a wide spectrum of cell/tissue types provide an unprecedented opportunity to predict a map of CRMs and constituent TFBSs in the human genome. Many computational methods have been developed to explore these data individually or jointly[30]. For instance, as the large number of binding-peaks in a typical TF ChIP-seq dataset dwarfs earlier motif-finding tools (e.g., MEME[31] and BioProspector[32]) to find TFBSs of the ChIP-ed TF, new tools (e.g., DREME[33], MEME-ChIP[34], XXmotif [35] and Homer[36]) have been developed. However, some of these tools (e.g. MEME-ChIP) were designed to find primary motifs of the ChIP-ed TF in short sequences (∼200bp) around the binding-peak summits in a small number of selected binding peaks in a dataset due to their slow speed. Some faster tools (e.g. Homer, DREME, and XXmotif) are based on the discriminative motif-finding schema[37] by finding overrepresented *k*-mers in a ChIP-seq dataset, but they often fail to identify TFBSs with subtle degeneracy. As TFBSs form CRMs for combinatory regulation in higher eukaryotes [1, 38], tools such as SpaMo [39], CPModule [40] and CCAT [41] have been developed to identify multiple closely located motifs as CRMs in a single ChIP-seq dataset. However, these tools cannot predict CRMs containing novel TFBSs, because they all depend on a library of known motifs (e.g., TRANSFAC [42] or JASPAR [43]) to scan for cooperative TFBSs in binding peaks. Due probably to the difficulty to find TFBS motifs in a mammalian TF ChIP-seq dataset that may contain tens of thousands of binding peaks, few efforts have been made to explore entire sets of an increasing number of TF ChIP-seq datasets to simultaneously predict CRMs and constituent TFBSs [44-47].

On the other hand, as a single histone mark is not a reliable CRM predictor, a great deal of efforts have been made to predict CRMs based on multiple histone marks and chromatin accessibility (CA) data from the same cell/tissue types using various machine-learning methods, including hidden Markov models[48], dynamic Bayesian networks[49], time-delay neural networks[50], random forest[51], and support vector machines (SVMs)[52]. Many enhancer databases have also been created either by combining results of multiple such methods[53-55], or by identifying overlapping regions of CA and histone mark tracks in the same cell/tissue types[56-60]. In particular, the ENCODE phase 3 consortium[26] recently identified 926,535 candidate *cis*-regulatory elements (cCREs) based on overlaps between millions of DNase I hypersensitivity sites (DHSs)[61] and transposase accessible sites (TASs)[62], active promoter histone mark H3K4me3[63] peaks, active enhancer mark H3K27ac[64] peaks, and insulator mark CTCT[65] peaks in a large number of cell/tissue types. Although CRMs predicted by these methods are often cell/tissue type-specific, their applications are limited to cell/tissue types for which the required datasets are available[26, 48, 49, 66]. The resolution of these methods is also low[48, 49, 66] and often lacks TFBSs information[26, 48, 49, 66], particularly for novel motifs, although some predictions provide TFBSs locations by finding matches to known motifs[54, 55, 59]. Moreover, results of these methods are often inconsistent[67-70], e.g., even the best-performing tools (DEEP and CSI-ANN) have only 49.8% and 45.2%, respectively, of their predicted CRMs overlap with the DHSs in Hela cells[52]; and only 26% of predicted ENCODE enhancers in K562 cells can be experimentally verified[67]. The low accuracy of these methods might be due to the fact that CA and histone marks alone are not reliable predictors of active CRMs [52, 67, 68, 70].

It has been shown that TF binding data are more reliable for predicting CRMs than CA and histone mark data, particularly, when multiple closely located binding sites for key TFs were used [52, 67, 68, 70]. Moreover, although primary binding sites of a ChIP-ed TF tended to be enriched around the summits of binding peaks, TFBSs of cooperators of the ChIP-ed TFs tend to appear at the two ends of binding peaks[71, 72]. With this recognition, instead of predicting cell/tissue type specific CRMs using CA and histone marks data, we proposed to first predict a largely cell-type agnostic or static map of CRMs and constituent TFBBs in the genome by integrating all available TF ChIP-seq datasets for different TFs in various cell/tissue types[46, 47], just as has been done for identifying all genes encoded in the genome using gene expression data from all cell/tissue types[73]. We also proposed to appropriately extend short binding peaks to the typical length of enhancers, so that more TFBSs for cooperators of the ChIP-ed TF could be included [71, 72], and thus, full-length CRMs could be identified[46, 47]. Once a map of CRMs and constituent TFBSs is available, the specificity of CRMs in any cell/tissue type can be determined using one or few epigenetic mark datasets collected in the cell/tissue type[26], because when anchored by correctly predicted CRMs, the accuracy of epigenetic marks for predicting active CRMs could be largely improved [68]. Although our earlier implementation of this strategy, dePCRM, resulted in promising results using even insufficient datasets available then[46, 47], we were limited by three technical hurdles. First, although existing motif-finders such as DREME used in dePCRM worked well for relatively small ChIP-seq datasets from organisms with smaller genomes such as the fly [47], they are unable to work on very large entire datasets from mammalian cells/tissues, so we had to split a large dataset into smaller ones for motif finding in the entire dataset[46], which may compromise the accuracy of motif finding and complicate subsequent data integration. Second, although the distances and interactions between TFBSs in a CRM are critical, both were not considered in our earlier scoring functions [46, 47], potentially limiting the accuracy of predicted CRMs. Third, the earlier “branch-and-bound” approach to integrate motifs found in different datasets is not efficient enough to handle a much larger number of motifs found in an ever increasing number of large ChIP-seq datasets from human cells/tissues[46, 47]. To overcome these hurdles, we developed dePCRM2 based on an ultrafast, accurate motif-finder ProSampler[72], a novel effective combinatory motif pattern discovery method, and scoring functions that model essentials of both the enhanceosome and billboard models of CRMs[74-76]. Using available 6,092 ChIP-seq datasets covering 77.47% of the human genome after extending the binding peaks, dePCRM2 was able to partition the covered genome regions into a CRM candidate (CRMC) set and a non-CRMC set, and predict 201 unique TF binding motif families in the CRMCs. Both evolutionary and independent experimental data indicate that at least the vast majority of the predicted 1,404,973 CRMCs might be functional, while at least the vast majority of the predicted non-CRMCs might not be functional.

## Results

### The dePCRM2 pipeline

TFs in higher eukaryotes tend to cooperatively bind to their TFBSs in CRMs[1]. Different CRMs of the same gene are structurally similar and closely located[77]. For example, in the locus control region (LCR) of the hemoglobin genes in the mouse genome, multiple enhancers with similar combinations of TFBSs regulate the expression of different hemoglobin genes in different tissues and developmental stages [78]. Moreover, functionally related genes are often regulated by the same sets of TFs in different cell types during development and in maintaining physiological homeostasis[1]. Due to the clustering nature of TFBSs of cooperative TFs in a CRM, if we extend the called short binding peaks of a TF ChIP-seq dataset from the two ends and reach the typical size of a CRM (500∼3,000bp)[79], the extended peaks would have a great chance to contain TFBSs of cooperative TFs[46, 47, 72]. For instance, if two different TFs cooperatively regulate the same regulons in several cell types, then at least some of the extended peaks of datasets for the two TFs from these cell types should contain the TFBSs of both TFs, and even have some overlaps if the CRMs are reused in different cell types. Therefore, if we have a sufficient number of ChIP-seq datasets for different TFs from the same and different cell types, we are likely to include datasets for some cooperative TFs, and their TFBSs may co-occur in some extended peaks. Based on these observations, we designed dePCRM[46, 47] and dePCRM2 to predict CRMs and constituent TFBSs by identifying overrepresented co-occurring patterns of motifs found by a motif-finder in a large number of TF ChIP-seq datasets. dePCRM2 overcomes the aforementioned shortcomings of dePCRM as follows. First, using an ultrafast and accurate motif-finder ProSampler[72], we can find significant motifs in available ChIP-seq datasets of any size (Figures 1A and 1B) without the need to split large datasets into small ones[46]. Second, after identifying highly co-occurring motifs pairs (CPs) in the extended binding peaks in each dataset (Figure 1C), we cluster highly similar motifs in the CPs and find a unique motif (UM) in each resulting cluster (Figure 1D). Third, we model distances and interactions among cognate TFs of the binding sites in a CRM by constructing interaction networks of the UMs based on the distance between the binding sites and the extent to which biding sites in the UMs cooccur to improve prediction accuracy (Figure 1E). Fourth, we identify as CRMCs closely located clusters of binding sites of the UMs along the genome (Figure 1F), thereby partitioning genome regions covered by the extended binding peaks into a CRMCs set and a non-CRMCs set. Fifth, we evaluate each CRMC using a novel score that considers not only the number of TFBSs in a CRM, but also the distances between the TFBSs, their quality scores and all pair-wise cooccurring frequencies between their motifs (Figure 1G). Lastly, we compute a p-value for each *S*_*CRM*_ score, so that CRMs and constituent TFBSs can be predicted at different significant levels using different *S*_*CRM*_ score or p-value cutoffs. Clearly, as the number of UMs is a small constant number constrained by the number of TF families encoded in the genome, the downstream computation based on the set of UMs runs in a constant time, thus dePCRM2 is highly scalable. The source code of dePCRM2 is available at http://github.com/zhengchangsulab/pcrm2

**Figure 1.**
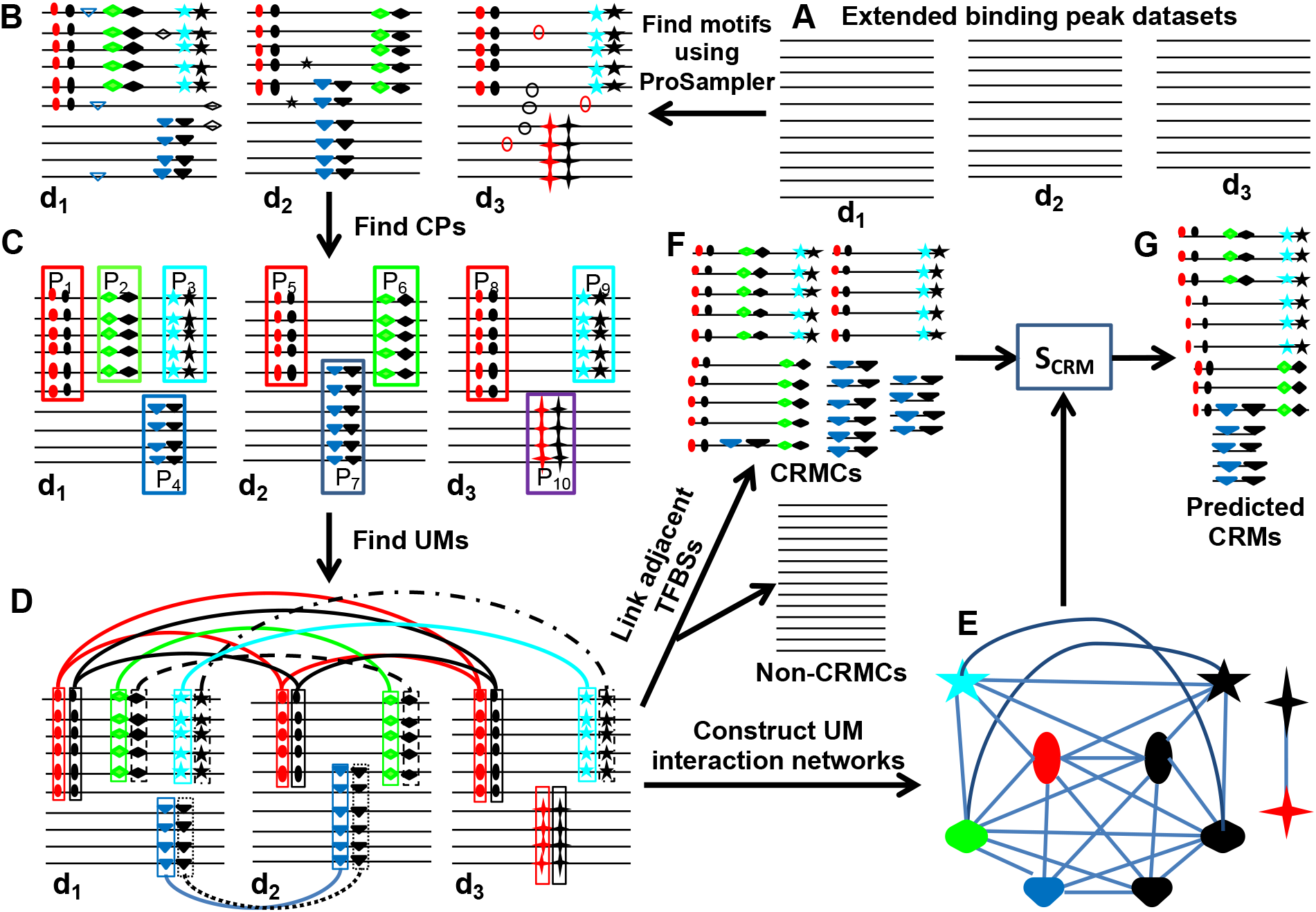
Schematic of the dePCRM2 pipeline. A. Extend each binding peak in each dataset to its two ends to reach a preset length, i.e., 1,000bp. B. Find motifs in each dataset using ProSampler. C. Find CPs in each dataset. For clarity, only the indicated CPs are shown, while those formed between motifs in pairs P_1_ and P_2_ in dataset d_1_, and so on, are omitted. D. Construct the motif similarity graph, cluster similar motifs and find UMs in the resulting motif clusters. Each node in the graph represents a motif, while weights on the edges are omitted for clarity. Clusters are connected by edges of the same color and line type. E. Construct UM interaction networks. Each node in the networks represents a UM, while weights on the edges are omitted for clarity. F. Project binding sites in the UMs back to the genome and link adjacent TFBSs along the genome, thereby identifying CRMCs and non-CRMCs. G. Evaluate each CRMC by computing its *S*_*CRM*_ score and the associated p-value.

### Unique motifs recall most known TF motifs families and have distinct patterns of interactions

ProSampler identified at least one motif in 5,991 (98.70%) of the 6092 ChIP-seq datasets (Supplementary Note) but failed to find any motifs in the remaining 101 (1.66%) datasets that all contain less than 310 binding peaks (Table S1), indicating that they are likely of low quality. As shown in Figure 2A, the number of motifs found in a dataset generally increases with the increase in the number of binding peaks in the dataset, but enters a saturation phase and stabilizes around 250 motifs when the number of binding peaks is beyond 40,000. In total, ProSampler identified 856,793 motifs in the 5,991 datasets. dePCRM2 found co-occurring motif pairs (CPs) in each dataset (Figure 1C) by computing a cooccurring score S_c_ for each pair of motifs in the dataset (formula 2). As shown in Figure 2B, *S*_*c*_ scores show a trimodal distribution. dePCRM2 selected as CPs the motif pairs that accounted for the mode with the highest S_c_ scores, and discarded those that accounted for the other two modes with lower S_c_ scores, because these low-scoring motif pairs were likely to co-occur by chance. In total, dePCRM2 identified 4,455,838 CPs containing 226,355 (26.4%) motifs from 5,578 (93.11%) of the 5,991 datasets. Therefore, we further filtered out 413 (6.89%) of the 5,991 datasets because each had a low S_c_ score compared with other datasets. Clearly, more and less biased datasets are needed to rescue their use in the future for more complete predictions. Clustering the 226,355 motifs in the CPs resulted in 245 clusters consisting of 2∼72,849 motifs, most of which form a complete similarity graph or clique, indicating that member motifs in a cluster are highly similar to each other (Figure S1A). dePCRM2 found a UM in 201 (82.04%) of the 245 clusters (Figure S1B and Table S1) but failed to do so in 44 clusters due to the low similarity between some member motifs (Figure S1A). Binding sites of the 201 UMs were found in 39.87∼100% of the sequences in the corresponding clusters, and in only 1.49% of the clusters binding sites were not found in more than 50% of the sequences due to the low quality of member motifs (Figure S2). Thus, this step retained most of putative binding sites in most clusters. The UMs contain highly varying numbers of binding sites ranging from 64 to 13,672,868 with a mean of 905,288 (Figure 2C and Table S1), reminiscent of highly varying number of binding peaks in the datasets (Supplementary Note). The lengths of the UMs range from 10 to 21bp with a mean of 11bp (Figure 2D), which are in the range of the lengths of known TF binding motifs, although they are biased to 10bp due to the limitation of the motif-finder to find longer motifs. As expected, a UM is highly similar to its member motifs that are highly similar to each other (Figure S1A). For example, UM44 contains 250 highly similar member motifs (Figure 2E). Of the 201 UMs, 117 (58.2%) match at least one of the 856 annotated motifs in the HOCOMOCO [80] and JASPAR[81] databases, and 92 (78.63%) match at least two (Table S2), suggesting that most UMs might consist of motifs of different TFs of the same TF family/superfamily that recognize highly similar motifs, a well-known phenomenon[82, 83]. Thus, a UM might represent a motif family/superfamily for the cognate TF family/superfamily. For instance, UM44 matches known motifs of nine TFs of the “ETS” family ETV4∼7, ERG, ELF3, ELF5, ETS2 and FLI1, a known motif of NFAT5 of the “NFAT-related factor” family, and a known motif of ZNF41 of the “more than 3 adjacent zinc finger factors” family (Figure 2F and Table S2). The high similarity of these motifs suggest that they might form a superfamily. The remaining 84 (43.28%) of the 201 UMs might be novel motifs recognized by unknown TFs (Figure S1B and Table S1). On the other hand, 64 (71.91%) of the 89 annotated motif TF families match one of the 201 UMs (Table S3), thus, our predicted UMs include most of the known TF motif families.

**Figure 2.**
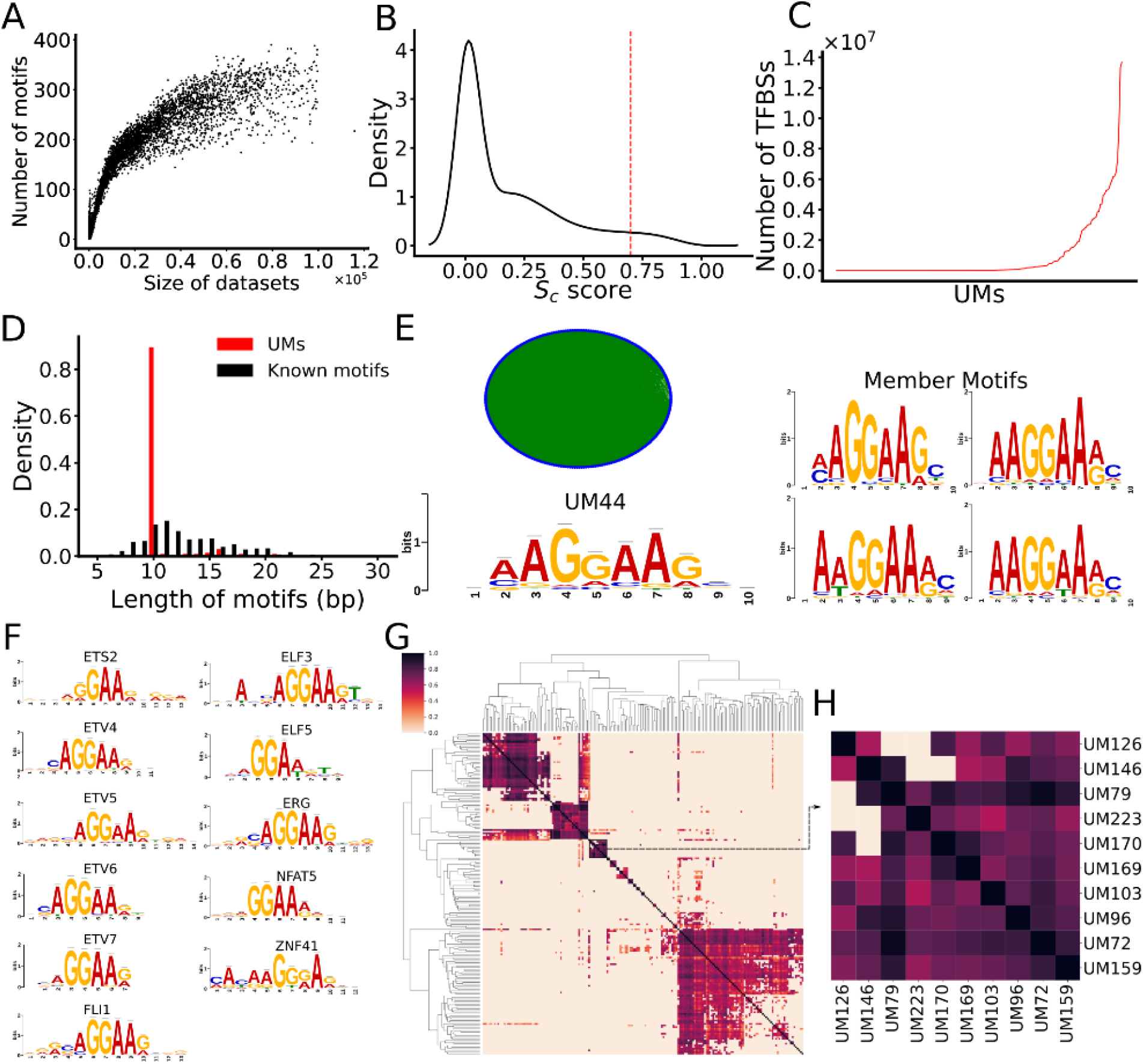
Prediction of UMs. A. Relationship between the number of predicted motifs in a dataset and the size of the dataset (number of binding peaks in the dataset). The datasets are sorted in ascending order of their sizes. B. Distribution of cooccurrence scores (*Sc*) of motif pairs found in a dataset. The dotted vertical line indicates the cutoff value (0.7) of *Sc* for predicting cooccurring pairs (CPs). C. Number of putative binding sites in each of the UMs sorted in ascending order. D. Distribution of the lengths of the UMs and known motifs in the HOCOMOCO and JASPAR databases. E. The logo and similarity graph of the 250 member motifs of UM44. In the graph, each node in blue represents a member motif, and two member motifs are connected by an edge in green if their similarity is greater than 0.8 (SPIC score). Four examples of member motifs are shown in the right panel. F. UM44 matches known motifs of nine TFs of the “ETS”, “NFAT-related factor”, and “more than 3 adjacent zinc finger factors” families. G. Heatmap of the interaction networks of the 201 UMs, while names of the UMs are omitted for clarity. H. A blowup view of the indicated cluster in G, formed by 10 UMs, of which UM126, UM146, UM79, UM223, UM170, UM103 and UM159 match known motifs of MESP1|ZEB1, TAL1::TCF3, ZNF740, MEIS1|TGIF1|MEIS2|MEIS3, TCF4|ZEB1|CTCFL|ZIC1|ZIC4|SNAI1, GLI2|GLI3 and KLF8, respectively. Some of these TFs are known collaborators in transcriptional regulation.

To model interactions between cognate TFs of the UMs, we computed an interaction score *S*_*INTER*_ based on distances and cooccurrence levels between binding sites of two UMs (formula 3), which largely improves our earlier score (data not shown) that only considers cooccurring frequencies of binding sites in two motifs [46, 47]. As shown in Figure 2G, there are clear interaction patterns between putative cognate TFs of many UMs, many of which are supported by experimental evidence. For example, in a cluster formed by 10 UMs (Figure 2H), seven of them (UM126, UM146, UM79, UM223, UM170, UM103 and UM159) match known motifs of MESP1/ZEB1, TAL1::TCF3, ZNF740, MEIS1/TGIF1/MEIS2/MEIS3, TCF4/ZEB1/CTCFL/ZIC1/ZIC4/SNAI1, GLI2/GLI3 and KLF8, respectively. At least a few of them are known collaborators in transcriptional regulation. For example, GLI2 cooperates with ZEB1 to repress the expression of *CDH*1 in human melanoma cells via directly binding to two close binding sites at the *CDH*1 promoter[84]; ZIC and GLI cooperatively regulate neural and skeletal development through physical interactions between their zinc finger domains [85]; and ZEB1 and TCF4 reciprocally modulate their transcriptional activities to regulate the expression of *WNT*[86], to name a few.

### An appropriate extension of original binding peaks greatly increases the power of datasets

By concatenating closely located binding sites of the UMs along the genome, dePCRM2 partitioned the 77.47% of the genome that are covered by the extended binding peaks (Supplementary Note) in two exclusive sets (Figure 1F), i.e., the CRMC set containing 1,404,973 CRMCs with a total length of bp (56.84%) covering 44.03% of the genome, and the non-CRMC set containing 1,957,936 sequence segments with a total length of 1,032,664,424bp (43.16%) covering 33.44% of the genome. Interestingly, only 57.88% (776,999,862bp) of genome positions of the CRMCs overlap those of the original binding peaks. Hence, dePCRM2 only retained 61.40% of genome positions covered by the original peaks, and abandoned the remaining 38.60% of nucleotide position. These abandoned positions covered by originally called binding peaks might not enrich for TFBSs, which is in agreement with earlier findings about the noisy nature of TF ChIP-seq data [87-89]. On the other hand, the remaining 42.12% (565,448,583bp) genome positions of the CRMCs overlap those of the extended parts of the original peaks, indicating that TFBSs of cooperative TFs are indeed enriched in the extended parts as has been shown earlier[46, 47, 71, 72], and dePCRM2 is able predict CRMs that are not covered by any binding peaks. Thus, by appropriately extending original binding peaks, we could greatly increase the power of datasets. Based on the overlap between a CRMC and original binding peaks in a cell/tissue type (Materials and Methods), dePCRM2 predicted functional states of the 57.88% of the CRMCs in at least one of the cell/tissue types from which binding peaks were called. However, dePCRM2 was not able to predict the functional states of the remaining 42.12% of the CRMCs that do not overlap any original binding peaks in the datasets. The predicted CRMCs and constituent TFBSs are available at https://cci-bioinfo.uncc.edu/

### The CRMCs are unlikely predicted by chance

To further evaluate the predicted CRMCs, we computed a *S*_*CRM*_ score for each CRMC (formula 4). As shown in Figure 3A, the distribution of the *S*_*CRM*_ scores of the CRMCs is strongly right-skewed relative to that of the Null CRMCs (Materials and Methods), indicating that the CRMCs generally score much higher than the Null CRMCs, and thus are unlikely produced by chance. Based on the distribution of the *S*_*CRM*_ scores of the Null CRMCs, dePCRM2 computed a p-value for each CRMC (Figure 3A). With the increase in the *S*_*CRM*_ cutoff α (*S*_*CRM*_ ≥ α), the associated p-value cutoff drops rapidly, while both the number of predicted CRMs and the proportion of the genome covered by the predicted CRMs decrease slowly (Figure 3B), indicating that most CRMCs have low p-values. For instance, with α increasing from 56 to 922, p-value drops precipitously from 0.05 to 1.00×10^−6^ (5×10^5^ fold), while the number of predicted CRMs decreases from 1,155,151 to 327,396 (3.53 fold), and the proportion of the genome covered by the predicted CRMs decreases from 43.47% to 27.82% (1.56 fold) (Figure 3B). Predicted CRMs contain from 20,835,542 (p-value ≤ 1×10^−6^) to 31,811,310 (p-value ≤ 0.05) non-overlapping putative TFBSs that consist of from 11.47% (p-value≤ 1×10^−6^) to 16.54% (p-value ≤ 0.05) of the genome (Figure 3C). In other words, dependent on p-value cutoffs (1×10^−6^ ∼0.05), 38.05∼41.23% of nucleotide positions of the predicted CRMs are made of putative TFBSs (Figure 3C), and most of predicted CRMs (93.99∼95.46%) and constituent TFBSs (93.20∼94.67%) are located in non-exonic sequences (NESs) (Figure 3C), comprising 26.66∼42.47% and 10.94∼16.03% of NESs, respectively (Figure 3D). Surprisingly, dependent on p-value cutoffs (1×10^−6^ ∼0.05), the remaining 4.54∼6.01% and 5.33∼6.80% of the predicted CRMs and constituent TFBSs, respectively, are in exonic sequences (ESs, including CDSs, 5’- and 3’-untranslated regions), respectively (Figure 3C), in agreement with an earlier report[90].

**Figure 3.**
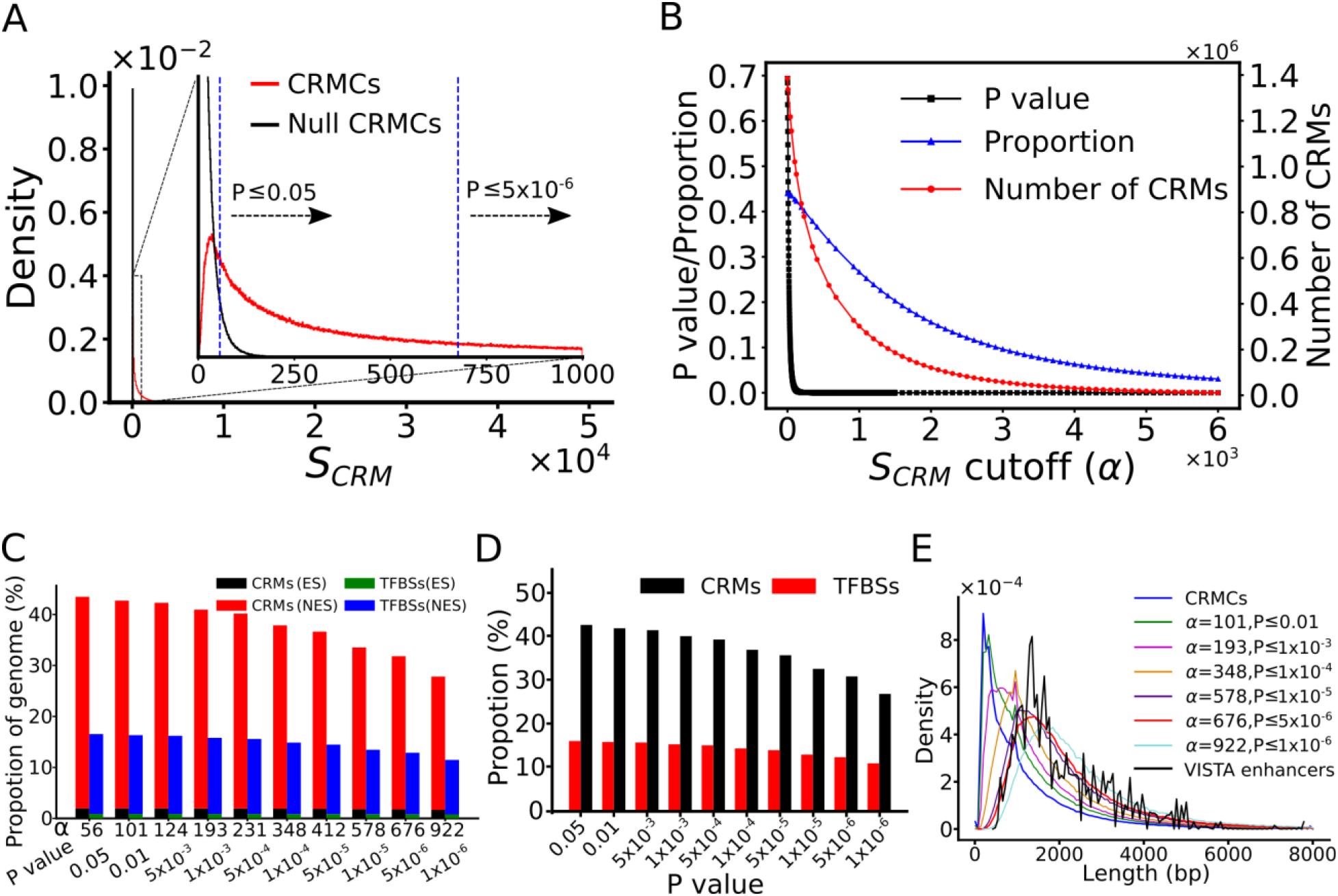
Prediction of CRMs using different *S* _*CRM*_ cutoffs. A. Distribution of *S* _*CRM*_ scores of the CRMCs and Null CRMCs. The inset is a blowup view of the indicated region. The vertical dashed lines indicate the associated p-values of the *S* _*CRM*_ cutoffs mentioned in the main text. B. Number of the predicted CRMs, proportion of the genome predicted to be CRMs and the associated p-value as functions of the S_CRM_ cutoff α. C. Percentage of the genome that are predicted to be CRMs and TFBSs in ESs and NESs using various S_CRM_ cutoffs and associated p-values. D. Percentage of NESs that are predicted to be CRMs and TFBSs using various S_CRM_ cutoffs and associated p-values. E. Distribution of the lengths of CRMs predicted using different S_CRM_ cutoffs and associated p-values.

### The *S*_*CRM*_ score captures the length feature of enhancers

As shown in Figure 3E, the CRMCs with a mean length of 981bp are generally shorter than VISTA enhancers with a mean length of 2,049bp. Specifically, 621,842 (44.26%) of the 1,404,973 CRMCs are shorter than the shortest VISTA enhancer (428bp), suggesting that they might be short CRMs (such as promoters or short enhancers) or components of long CRMs. However, these shorter CRMCs (< 428bp) comprise only 7.42% of the total length of the CRMCs. The remaining 733,132 (55.74%) CRMCs comprising 92.58% of the total length of the CRMCs are longer than the shortest VISTA enhancer (428bp), thus most of them are likely full-length CRMs. Therefore, predicted CRMC positions in the genome are mainly covered by full-length or longer CRMCs. As expected, with the increase in α (decrease in p-value cutoff), the distribution of the lengths of the predicted CRMs shifts to right and even surpass that for VISTA enhancers (Figure 3E), indicating shorter CRMCs can be effectively filtered out by a higher *S*_*CRM*_ cutoff α (a smaller p-value). The remaining CRMCs might be different type of CRMs with different length features. For instance, at a rather stringent *S*_*CRM*_ cutoff α =676 (p=5×10^−6^), 976,345 (69.49%) shorter CRMCs with a mean length of 387bp were filtered out (Figure 3E), the remaining 428,628 (30.51%) CRMCs have similar length distribution (mean length of 2292bp) to that of VISTA enhancers (mean length of 2049bp) (Figure 3E), which are mainly involved in development-related functions and are generally longer than other types of enhancers [91]. However, it is worth noting that VISTA enhancers may not necessarily all be in their full-length forms, because even a portion of an enhancer could be still partially functional[1], and it is still technically difficult to validate very long enhancers in transgene animal models in a large scale. Therefore, it is not surprising that with even more stringent *S*_*CRM*_ cutoffs, the predicted CRMs could be longer than VISTA enhancers (Figure 3C), and they are likely super-enhancers for cell differentiation of development[92]. Taken together, these results suggest that the *S*_*CRM*_ score captures the length feature of enhancers.

### The CRMCs and non-CRMCs show dramatically distinct evolutionary behaviors

To see how effectively dePCRM2 partitions the covered genome regions into the CRMC set and the non-CRMC set, we compared their evolutionary behaviors with those of the entire set of VISTA enhancers using the GERP[93] and phyloP[94] scores of their nucleotide positions in the genome. Both the GERP and the phyloP scores quantify conservation levels of genome positions based on nucleotide substitutions in alignments of multiple vertebrate genomes. The larger a positive GERP or phyloP score of a position, the more likely it is under negative/purifying selection; and a GERP or phyloP score around zero means that the position is selectively neutral or nearly so[93, 94]. However, a negative GERP or phyloP score is cautiously related to positive selection[93, 94]. For convenience of discussion, we consider a position with a GERP or phyloP score within an interval centering on 0 [-*δ*,+ *δ*] (*δ* >0) to be selectively neutral or nearly so, and a position with a score greater than *δ* to be under negative selection. We define proportion of neutrality of a set of positions to be the size of the area under the density curve of the distribution of the scores of the positions within the window [-*δ*,+ *δ*]. Because ESs evolve quite differently from NESs, we focused on the CRMCs and constituent TFBSs in NESs, and left those that overlap ESs in another analysis (Jing Chen, Pengyu Ni, Jun-tao Guo and Zhengchang Su). The choice of *δ* = 0.1, 0.2, 0.3, 0.4, 0.5, 1 and 2 gave similar results (data not shown), so we choose *δ* =1 in the subsequent analyses. As shown in Figure 4A, GERP scores of VISTA enhancers show a trimodal distribution with a small peak around score −5, a blunt peak around score 0, a sharp peak around score 3.5, and a small proportion of neutrality of 0.23, indicating that most nucleotide positions of VISTA enhancers are under strong evolutionary selection, particularly, negative selection. This result is consistent with the fact that VISTAT enhancers are mostly ultra-conserved[95], development-related enhancers[96, 97]. The 0.23 proportion of neutrality of the VISTA enhancer positions indicates that this proportion of positions might simply serve as non-functional spacers between adjacent TFBSs. Interestingly, there are 942 genome regions in the VISTA database, which failed to be validated as active enhancers in transgenic assays, and we found that they had similar GERP and phyloP score distributions as VISTA enhancers, although the former set is slightly less conserved than the latter set (Figure S3), suggesting that most of these “validated negative regions (VNRs) might actually have *cis*-regulatory functions under conditions that might have not be tested. In contrast, the distribution of the GERP scores of the non-CRMCs (1,034,985,426 bp) in NESs displays a sharp peak around score 0, with low right and left shoulders, and a high proportion of neutrality of 0.71 (Figure 4A), suggesting that most positions of the non-CRMCs are selectively neutral or nearly so, and thus are likely to be nonfunctional. The remaining 0.29 portion of positions of the non-CRMCs seem to be under varying levels of selection (Figure 4A), so they might have other functions than cis-regulation. Intriguingly, the distribution of the GERP scores of the 1,292,356 CRMCs (1,298,719,954bp) in NESs has a blunt peak around score 0, with high right and left shoulders, and a small proportion of neutrality of 0.31 (Figure 4A). Thus, like VISTA enhancers, most positions of the CRMCs are also under strong evolutionary selections, and thus, are likely to be functional, while the small proportion (0.31) of neutrality indicates that this proportion of positions in the CRMCs might simply serve as non-functional spacers, instead of TFBSs. Notably, the distribution of GERP scores of the CRMCs lack obvious peaks around scores −5 and 3.5 (Figure 4A), therefore, the average selection strength on the CRMCs is weaker than that on VISTA enhancers (but see the section “The higher the S_CRM_ score of a CRMC, the stronger evolutionary constraint it is under”). Nonetheless, this is expected considering the ultra-conversation nature of the small set of development-related VASTA enhancers[95-97]. In any rate, the dramatic differences between the evolutionary behaviors of the non-CRMCs and those of the CRMCs strongly suggests that dePCRM2 largely partitions the covered genome regions into a cis-regulatory CRMC set and a non-cis-regulatory non-CRMC set. Similar results were obtained using the phyloP scores, although they display quite different distributions than the GERP scores (Figure S4A).

**Figure 4.**
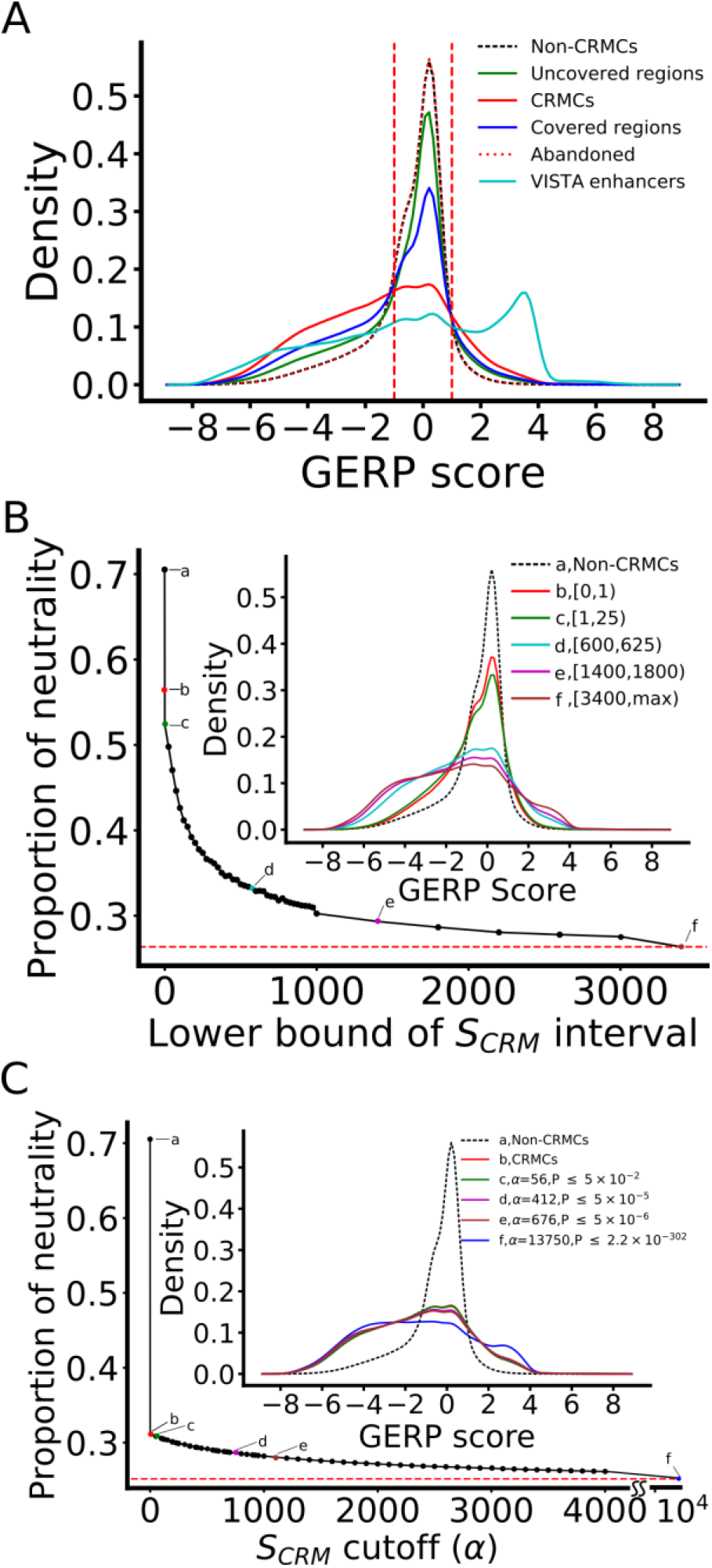
CRMCs and non-CRMCs in NESs show different evolutionary behaviors measured by GERP scores. A. Distributions of the GERP scores of nucleotide positions of VISTA enhancers, CRMCs, non-CRMCs, abandoned genome regions covered by original binding peaks, genome regions covered by extended binding peaks and genome regions uncovered by extended binding peaks. The area under the density curves in the score interval [-1, 1] is defined as the proportion of neutrality of the sequences. B. Proportion of neutrality of CRMCs with a S_CRM_ score in different intervals in comparison with that of the non-CRMCs (a). The inset shows the distributions of the GERP scores of the non-CRMCs and CRMCs with S_CRM_ scores in the intervals indicted by color and letters. C. Proportion of neutrality of CRMs predicted using different S_CRM_ score cutoffs and associated p-values in comparison with those of the non-CRMCs (a) and CRMCs (b). The inset shows the distributions of the GERP scores of the non-CRMCs, CRMCs and the predicted CRMs using the S_CRM_ score cutoffs and p-values indicated by color and letters. The dashed lines in B and C indicate the saturation levels.

To see why dePCRM2 abandoned the 38.60% nucleotide positions covered by the original binding peaks in predicting the CRMCs, we plotted the distribution of their conservation scores. As shown in Figure 4A, these abandoned positions have a GERP score distribution almost identical to those in the non-CRMCs, indicating that, like the non-CRMCs, they are largely selectively neutral, and thus, unlikely to be cis-regulatory, strengthening our earlier argument that they might not contain TFBSs. Therefore, dePCRM2 is able to accurately distinguish cis-regulatory and non-cis-regulatory parts in both the original binding peaks and their extended parts. As shown in Supplementary Note, the 10 CRM function-related elements datasets (Tables S4∼S8) that we collected for validating the predicted CRMCs are strongly biased to the covered genome regions relative to the uncovered regions. To see why this is possible, we plotted the distributions of conservation scores of the positions of the covered and uncovered regions in NESs. Interestingly, the uncovered regions have a GERP score distribution and a proportion of neutrality (0.59) in between those of the covered regions (0.49) and those of the non-CRMCs (0.71) (Figure 4A), indicating that the uncovered regions are more evolutionarily selected than the non-CRMCs as expected, but less evolutionary selected than the covered regions. This implies that the uncovered regions contain functional elements such as CRMs, but their density could be lower than that of the covered regions. Assuming that the total length of CRMs in a region is proportional to the total length of evolutionarily constrained parts in the region, the proportion of uncovered regions that might be CRMs could be estimated to be (1-0.59)/(1-0.49)=80.04% of that in the covered regions. Therefore, it appears that existing studies are strongly biased to more evolutionary constrained regions due probably to their large effect sizes and more critical functions. Similar results were obtained using the phyloP scores (Figure S4A).

### The higher the S_CRM_ score of a CRMC, the stronger evolutionary constraint it is under

To see whether the *S*_*CRM*_ score of a CRMC captures the strength of evolutionary selection that it is under, we plotted the distributions of the conservation scores of subsets of the CRMCs with a *S*_*CRM*_ score in different non-overlapping intervals. Remarkably, even the subset with *S*_*CRM*_ scores in the lowest interval [0, 1) has a smaller proportion of neutrality (0.56) than the non-CRMCs (0.71) (Figure 4B), indicating that even these low-scoring CRMCs with short lengths (Figure 3E) are more likely to be under strong evolutionary constraints than the non-CRMCs, and thus might be more likely cis-regulatory. With the increase in the lower bound of *S*_*CRM*_ intervals, the proportion of neutrality of the corresponding subsets of CRMCS drops rapidly, followed by a slow linear decrease around the interval [1000,1400) (Figure 4B). Therefore, the higher the *S*_*CRM*_ score of a CRMC, the more likely it is under strong evolutionary constraint, suggesting that the *S*_*CRM*_ score indeed captures the evolutionary behavior of a CRM as a functional element, in addition to its length feature (Figure 3E). The same conclusion can be drawn from the phyloP scores (Figure S4B).

We next examined the relationship between the conservation scores of the predicted CRMs and *S*_*CRM*_ score cutoffs α (or p-value cutoffs) used for their predictions. As shown in Figure 4C, even the CRMs predicted at a low α have a much smaller proportion of neutrality (e.g., 0.31 for the smallest α=0, i.e., the entire CRMC set) than the non-CRMCs (0.71), suggesting that most of the predicted CRMs might be authentic although some short ones may not be in full-length, while the non-CRMCs might contain few false negative CRMCs. With the increase in α (decrease in p-value cutoff), the proportion of neutrality of the predicted CRMs decreases but slowly, entering a saturation phase (Figure 4C). Interestingly, at very high *S*_*CRM*_ score cutoffs, the predicted CRMs evolve like VISTA enhancers, with a trimodal GERP score distribution, and thus might be involved in development[98, 99]. For instance, at α= 13,750, the distribution of GERP scores of the predicted CRMs displays a peak around score −5 and a peak around score 3.5, with a small proportion of neutrality of 0.24 (Figure 4C) (it is 0.23 for VISTA enhancers, Figure 4A). Thus, the higher α (i.e., the smaller the p-value cutoff), the more likely the predicted CRMs are under strong evolutionary constraints. The infinitesimal decrease in the proportion of neutrality of predicted CRMs with the increase in *S*_*CRM*_ cutoffs (Figure 4C) strongly suggests that the predicted CRMs, particularly those at a low p-value cutoff, are under similarly strong evolutionary constraints, close to the possibly highest saturation level to which ultra-conserved VISTA enhancers are subject. Therefore, it is highly likely that at low p-value cutoffs, specificity of the predicted CRMs might approach the possibly highest level that the VISTA enhancers achieve. However, without the availability of a gold standard negative CRM set in the genome[23], we could not explicitly calculate the specificity of the predicted CRMs at different p-value cutoffs. Similar results are observed using the phyloP scores (Figure S4C).

### dePCRM2 achieves high sensitivity and likely high specificity for recalling functionally validated CRMs and non-coding SNPs

To further evaluate the accuracy of dePCRM2, we calculated the sensitivity (recall rate or true positive rate (TPR)) of CRMs predicted at different *S*_*CRM*_ cutoffs α and associated p-values for recalling a variety of CRM function-related elements located in the covered genome regions in the 10 experimentally determined datasets in various cell/tissue types (Tables S4∼S8, Materials and Methods). Here, if a predicted CRM and an element overlap each other by at least 50% of the length of the shorter one, we say that the CRM recalls the element. As shown in Figure 5A, with the increase in the p-value cutoff, the sensitivity for recalling the elements in all the 10 datasets increases rapidly and becomes saturated well before p-value increases to 0.05 (α ≥ 56). Figures S5A∼S5J show examples of the predicted CRMs overlapping and recalling the elements in the 10 datasets. Particularly, at p-value cutoff 5×10^−5^ (α=412), the predicted 593,731 CRMs covering 36.63% of the genome (Figure 3C) recall 100% of VISTA enhancers[79] and 91.61% of ClinVar SNPs[79] (Figure 5A). The rapid saturation of sensitivity for recalling these two types of validated functional elements at such a low p-value cutoff once again strongly suggests that dePCRM2 also achieves very high specificity, although we could not explicitly compute it for the aforementioned reason. On the other hand, even at the higher p-value cutoff 0.05 (α=56), the predicted 1,155,151 CRMs covering 43.47% of the genome (Figure 3C) only achieve varying intermediate levels of sensitivity for recalling FANTOM5 promoters (FPs)(88.77%)[100], FANTOM5 enhancers (FEs) (81.90%)[101], DHSs (74.68%)[61], TASs (84.32%)[29], H3K27ac (82.96%)[29], H3K4me1 (76.77%)[29], H3K4me3 (86.96%)[29] and GWAS SNPs (64.50%)[102], although all are significantly higher than that (15%) of randomly selected sequences with matched lengths from the covered genome regions (Figure 5A).

**Figure 5.**
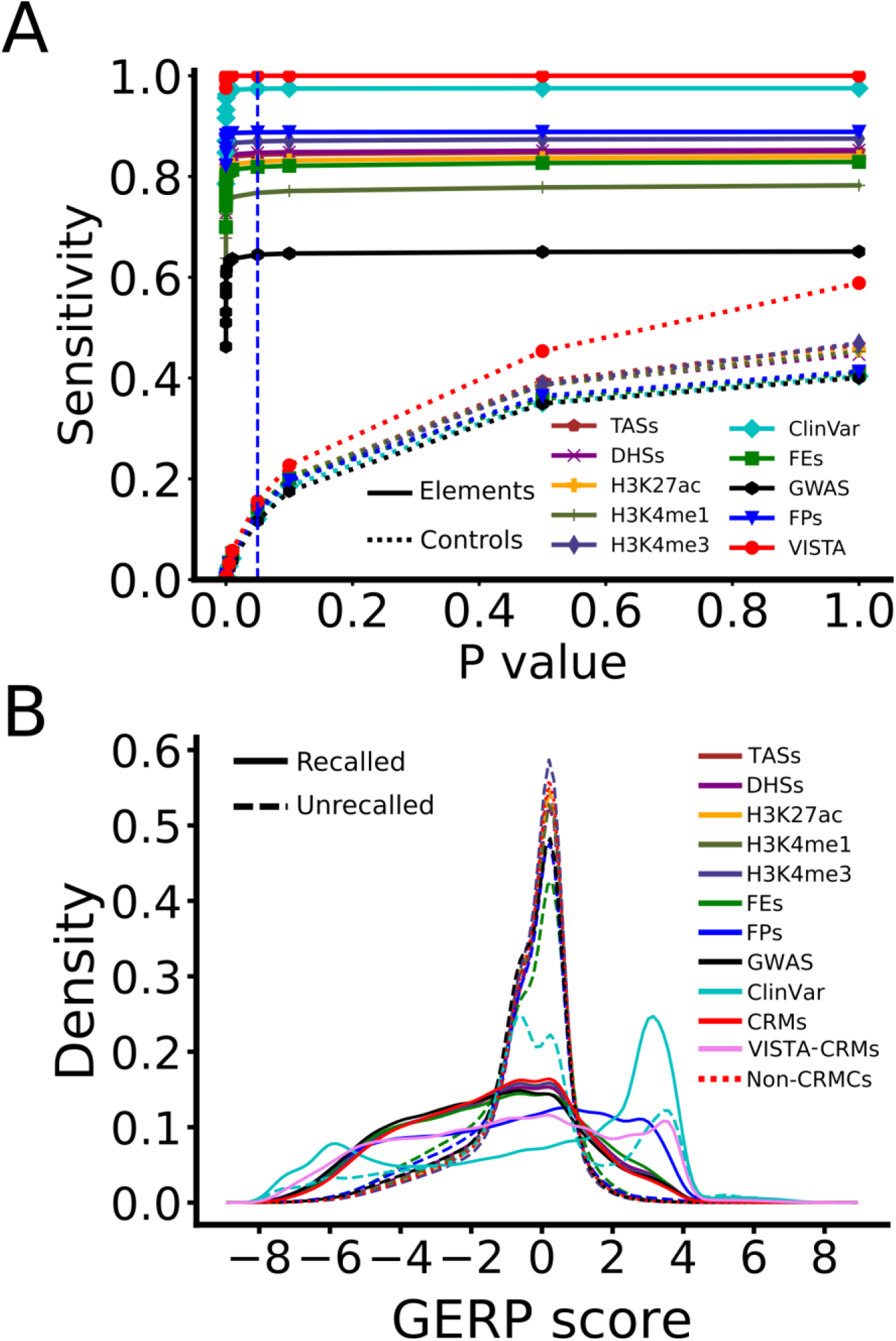
Validation of the predicted CRMs by 10 experimentally determined sequence elements datasets. A. Sensitivity (recall rate or TPR) of the predicted CRMs and control sequences as a function of p-value cutoff for recalling the sequence elements in the datasets. The dashed vertical lines indicate the p-value ≤0.05 cutoff. B. Distributions of GERP scores of the recalled and unrecalled elements in each dataset in comparison with those of the predicted CRMs at p≤0.05 and non-CRMCs. Note that there are no unrecalled VISTA enhancers, and the distribution of the recalled 785 VISTA enhancers in the covered genome regions (not shown) is almost identical to the entire set of 976 VISTA enhancers (Figure 4A). The curve labeled by VISTA-CRMs is the distribution of CRMs that overlap and recall the 785 VISTA enhancers.

To find out the reasons for such varying sensitivity of dePCRM2 for recalling different types elements in the 10 datasets, we plotted the distribution of GERP scores of the recalled and uncalled elements in each dataset by our predicted CRMs at p-value <0.05. Since we have already plotted the distribution of the entire set of VISTA enhancers (Figure 4A), to avoid redundancy, we instead plotted the distribution for the CRMs (VISTA-CRMs) that overlap and recall the 785 VISTA enhancers in the covered regions. As shown in Figure 5B, like the predicted CRMs, the recalled elements in all the datasets are under strong evolutionary selections (at p-value <0.05), thus are likely functional. However, VISTA-CRMs, recalled ClinVar SNPs and recalled FPs evolve more like VISTA enhancers with a trimodal GERP score distribution (Figure 4A), suggesting that they are under stronger evolution constraints than the other recalled element types. These results are not surprising, as we mentioned earlier VISTA enhancers are mostly ultra-conserved, development related enhancers[95-97], while ClinVar SNPs were identified for their conserved critical functions[103], and promoters are well-known to be more conserved than are enhancers[104]. In stark contrast, like the non-CRMCs, all unrecalled elements in the 10 datasets are largely selectively neutral, and thus, are unlikely to be functional, with the exception that the 10,350 (2.57%) unrecalled ClinVar SNPs display a trimodal distribution and there are no unrecalled VISTA enhancers (Figure 5B). Notably, proportions of neutrality of unrecalled PEs (0.59) and PFs(0.63) are smaller than that of the non-CRMCs (0.71) (Figure 5B), suggesting we might miss a small portion of authentic PEs and PFs (see below for false negative rate (FNR) estimations of our CRMs). Nevertheless, assuming that at least most of unrecalled elements in the datasets except the VISTA and ClinVar datasets, are non-*cis*-regulatory, we estimated that the false discovery rate (FDR) of the remaining eight datasets might be up to from 11.23% (1-0.8877) for FPs to 35.50% (1-0.6450) for GWAS SNPs. Such high FDRs for CA (DHSs and TASs) and histone marks are consistent with an earlier study[68]. Interestingly, the trimodal distribution of GERP scores of the 2.57% of unrecalled ClinVar SNPs displays a large peak around score 0 and two small peaks around −5 and 3.5, with a proportion of neutrality 0.40 (Figure 5B), indicating that about 40% of the relevant SNPs might be selectively neutral, and thus non-functional. We therefore estimated the FDR of the ClinVar SNP dataset to be about 0.40*2.57%=1.03%. Hence, like VISTA enhancers, ClinVar SNPs are a reliable set for evaluating CRM predictions. The peak of the unrecalled ClinVar SNPs around score 3.5 (Figure 5B), indicates that the relevant SNPs are under strong purifying selection, and thus might be functional, but were missed by dePCRM2. We therefore estimate our predictions (at p-value <0.05) might have a FNR < 2.57%-1.03%=1.54%. In other words, the real sensitivity (=1-FNR) for dePCRM2 to recall authentic ClinVar SNPs might be higher than the calculated 97.54% (Figure 5A). These estimates are supported by the zero FNR and 100% sensitivity for our predicted CRMs to recall VISTA enhancers (Figure 5A) and a simulation to be described later.

The zero, very low (<1.03%) and low (11.23%) FDRs of VISTA enhancers, ClinVar SNPs and FPs datasets, respectively, are clearly related to the high reliability of the experimental methods used to characterize them. However, the low FDRs might also be related to the highly conserved nature of these elements (Figure 5B), as their critical functions and large effect sizes may facilitate their correct characterization. In this regard, we note that the intermediately high FDRs of the FEs(18.10%), DHSs(25.32), TASs (15.68%), H3K4m3 (13.04%), H3K4m1 (23.23%) and H3K27ac (17.04%) datasets might be due to the facts that bidirectional transcription[105], CA[68, 70, 106] and histone marks[68, 70] are not unique to active enhancers. The very high FDR of GWAS SNPs (35.5%) might be due to the fact that a lead SNP associated with a trait may not necessarily be located in a CRM and causal; rather, some variants in a CRM, which are in LD with the lead SNP, are the culprits[102, 107]. Example of GWAS SNPs in LD with positions in a CRM are shown in Figures S5K and S5L. Interestingly, many recalled ClinVar SNPs (42.59%) and GWAS SNPs (38.18%) are located in critical positions in predicted binding sites of the UMs (e.g., Figures S5D and S5F).

In addition, we found that 722 (76.65%) of the 942 VNRs in the VISTA database fall in the covered 77.47% genome regions. At a p-value cutoff of 0.05, the predicted CRMs recall 711 (98.48%) of the 722 VNRs. Interestingly, recalled VNR positions evolve similarly to the VISTA enhancer positions, while unrecalled VNR positions evolve similarly to the non-CRMC positions (Figure S3). These results strongly suggest that recalled VNRs might be true enhancers that function in conditions yet to be tested, as acknowledged by the VISTA team [108]. On the other hand, most unrecalled VNRs might not be *cis*-regulatory.

### dePCRM2 outperforms state-of-the-art methods for predicting CRMs

We compared our predicted CRMs at p-value ≤ 0.05 (S_CRM_ < 56) with three most comprehensive sets of predicted enhancers/promoters, i.e., GeneHancer 4.14[55], EnhancerAtals2.0[59] and cCREs[26]. For convivence of discussion, we call these three sets enhancers or cCREs. GeneHancer 4.14 is the most updated version containing 394,086 non-overlapping enhancers covering 18.99% (586,582,674bp) of the genome (Figure 6A). These enhancers were predicted by integrating multiple sources of both predicted and experimentally determined CRMs, including VISTA enhancers[79], ENCODE phase 2 enhancer-like regions[109], ENSEMBL regulatory build[53], dbSUPER[110], EPDnew promoters[111], UCNEbase[112], CraniofacialAtlas[113], FPs[100] and FEs [101]. Enhancers from ENCODE phase 2 and ESEMBL were predicted based on multiple tracks of epigenetic marks using the well-regarded tools ChromHMM[48] and Segway[114]. Of the GeneHancer enhancers, 388,407 (98.56%) have at least one nucleotide located in the covered genome regions, covering 18.89% of the genome (Figure 6A). EnhancerAtlas 2.0 contains 7,433,367 overlapping cell/tissue-specific enhancers in 277 cell/tissue types, which were predicted by integrating 4,159 TF ChIP-seq, 1,580 histone mark, 1,113 DHS-seq, and 1,153 other enhancer function-related datasets, such as FEs[115]. After removing redundancy (identical enhancers in difference cell/tissues), we ended up with 3,452,739 EnhancerAtlas enhancers that may still have overlaps, covering 58.99% (1,821,795,020bp) of the genome (Figure 6A), and 3,417,629 (98.98%) of which have at least one nucleotide located in the covered genome regions, covering 58.78% (1,815,133,195bp) of the genome (Figure 6A). cCREs represents the most recent CRM prediction by the ENCODE phase 3 consortium[26], containing 926,535 non-overlapping cell type agnostic enhancers and promoters covering 8.20% (253,321,371bp) of the genome. The cCREs were predicted based on overlaps among 703 DHS, 46 TAS and 2,091 histone mark datasets in various cell/tissue types produced by ENCODE phases 2 and 3 as well as the Roadmap Epigenomics projects[26]. Of these cCREs, 917,618 (99.04%) have at least one nucleotide located in the covered genome regions, covering 8.13% (251,078,466bp) of the genome (Figure 6A). Thus, due probably to the aforenoted reasons, these three sets of predicted enhancers and cCREs also are strongly biased to the covered regions relative to the uncovered regions. Both the number (1,155,151) and genome coverage (43.47%) of our predicted CRMs (p-value<0.05) are larger than those of GeneHancer enhancers (388,407 and 18.89%) and of cCREs (917,618 and 8.12%), but smaller than those of EnhancerAtlas enhancers (3,417,629 and 58.78%), in the covered regions.

**Figure 6.**
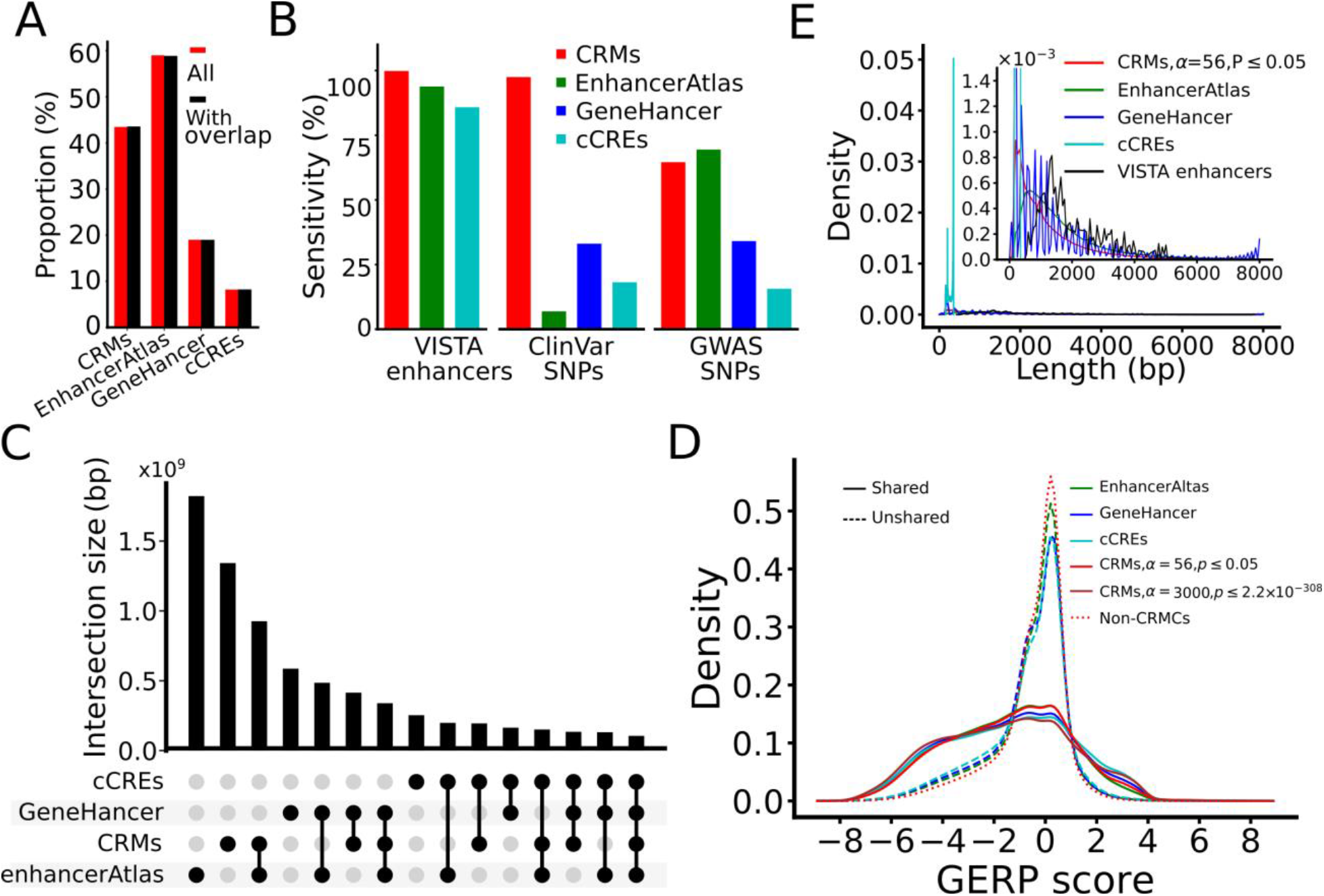
Comparison of the performance of dePCRM2 and three state-of-the-art methods. A. Proportion of the genome that are covered by enhancers/CRMs predicted by the four methods (All), and proportion of genome regions covered by predicted enhancers/CRMs that at least partially overlap the covered genome regions (With overlap). B. Sensitivity for recalling VISTA enhancers, ClinVar SNPs and GWAS SNPs, by the predicted enhancers/CRMs that at least partially overlap the covered genome regions. C. Upset plot showing numbers of nucleotide positions shared among the predicted CRMs, GeneHancer enhancers, EnhancerAtlas enhancers and cCREs. D. Distributions of GERP scores of nucleotide positions of the CRMs predicted at p-value ≤ 0.05 and p-value ≤ 2.2×10^−308^, and the non-CRMCs, as well as of nucleotide positions that GeneHancer enhancers, EnhancerAtlas enhancers and cCREs share and do not share with the predicted CRMs at p-value ≤ 0.05. E. Distributions of lengths of the four sets of predicted enhancers/CRMs in comparison to that of VISTA enhancers. The inset is a blow-up view of the axes defined region.

To make the comparisons fair, we first computed the sensitivity of these three sets of enhancers and cCREs for recalling VISTA enhancers, ClinVar SNPs and GWAS SNPs in the covered regions. We omitted FPs, FEs, DHSs, TASs and the three histone marks for the valuation as they were used in predicting CRMs by GeneHancer 4.14, EnhancerAtlas 2.0 or ENCODE phase 3 consortium. We also excluded VISTA enhancers for evaluating GeneHancer enhancers as the former were compiled in the latter [55]. Remarkably, our predicted CRMs outperform EnhancerAtlas enhancers for recalling VISTA enhancers (100.00% vs 94.01%) and ClinVar SNPs (97.43% vs 7.03%) (Figure 6B), even though our CRMs cover a smaller proportion of the genome (43.47% vs 58.78%, or 35.22% more) (Figure 6A), indicating that dePCRM2 has both higher sensitivity and specificity than the method behind EnhancerAtlas 2.0[59]. However, our CRMs underperform EnhancerAtlas enhancers for recalling GWAS SNPS (64.50% vs 69.36%, or 7.54% more) (Figure 6B). As we indicated earlier, the lower sensitivity of dePCRM2 for recalling GWAS SNPs might be due to the fact that an associated SNP may not necessarily be causal (Figures S5K and S5L). The higher sensitivity of EnhancerAtlas enhancers for recalling GWAS SNPs might be simply thanks to their 35.22% more coverage of the genome (58.78%) than that of our predicted CRMs (43.47%) (Figure 6A). Our predicted CRMs outperform cCREs for recalling VISTA enhancers (100% vs 85.99%), ClinVar SNPs (97.43% vs 18.28%) and GWAS SNPs (64.50% vs 15.74%) (Figure 6B). Our predicted CRMs also outperform GeneHancer enhancers for recalling ClinVar SNPs (97.43% vs 33.16%) and GWAS SNPs (64.50% vs 34.11%) (Figure 6B). However, no conclusion can be drawn from these results about the specificity of our predicted CRMs compared with GeneHancer enhancers and cCREs, because our predicted CRMs cover a higher proportion of the genome than both of them (43.47% vs 18.89% and 8.20%). On the other hand, both GeneHancer 4.14 enhancers and cCREs outperform EnhancerAtlas enhancers for recalling ClinVar SNPs (33.16% and 18.28% vs 7.03%)(Figure 6B), even though they have a much smaller genome coverage than EnhancerAtlas enhancers (18.89% and 8.20% vs 58.78%) (Figure 6A), indicating that they have higher specificity than EnhancerAtlas enhancers.

As shown in Figure 6C, the intersections/overlaps between the four predicted CRMs/enhancer/cCREs sets are quite low. For instance, EnhancerAtlas enhancers, GeneHancer enhancers and cCREs share 926,396,395bp (50.85%), 414,806,711bp (70.72%), and 194,709,825bp (76.86%) of their nucleotide positions with our predicted CRMs, corresponding to 69.01%, 30.90% and 14.51% of the positions of our CRMs (Figure 6C), respectively. There are only 105,606,214bp shared by all the four sets, corresponding to 5.80%, 18.00%, 41.69% and 7.87% of nucleotide positions covered by EnhancerAtlas enhancers, GeneHancer enhancers, cCREs and our CRMs, respectively. As expected, the 50.85%, 70.72% and 76.86% of their nucleotide positions that EnhancerAtlas enhancers, GeneHancer enhancers and cCREs share with our CRMs, respectively, evolve similarly to our predicted CRMs, although those of GeneHancer enhancers and cCREs are under slightly higher evolutionary constraints than our CRMs (Figure 6D). However, at a higher S_CRM_ cutoff, e.g. α=3,000 (p<2.2×10^−302^), our predicted CRMs are even under stronger evolutionary constraints than the shared GeneHancer enhancers and cCREs positions (Figure 6D). Therefore, the shared GeneHancer enhancers and cCREs positions just evolve like subsets of our predicted CRMs with higher S_CRM_ scores. By stark contrast, like the non-CRMCs, the remaining 49.14%, 29.28% and 23.13% of their nucleotide positions that EnhancerAtlas enhancers, GeneHancer enhancers and cCREs do not share with our CRMs, respectively, are largely selectively neutral, although they all have slightly smaller proportion of neutrality than that of the non-CRMCs (0.66, 0.63 and 0.61 vs. 0.71, respectively) (Figure 6D), due probably to the small FNR (<1.54%) of our predicted CRMs. Nonetheless, these results strongly suggest that the vast majority of the unshared positions of the three sets of predicted enhancers/eCREs are selectively neutral, and thus might be nonfunctional. It appears that the predicted enhancers/cCREs in the three sets that overlap our CRMs are likely to be authentic, while most of those that do not might be false positives. Hence, we estimated the FDR of EnhancerAtlas enhancers, GeneHancer enhancers and cCREs to be around 49.14%, 29.28% and 23.13%, respectively. Therefore, it is highly likely that GeneHancer 4.14 and cCREs might largely under-predict enhancers as evidenced the fact that they are targeted at evolutionarily more constrained elements (Figure 6D), even though they have rather high FDRs around 29.28% and 23.12%, respectively (Figure 6D), while EnhancerAtlas 2.0 might largely over-predict enhancers with a very high FPR around 49.14% (Figure 6D).

Finally, we compared the lengths of the four sets of predicted CRMs/enhancers/cCREs with those of VISTA enhancers. As shown in Figure 6E, the distribution of the lengths of cCREs has a narrow high peak at 345bp with a mean length of 273bp and a maximal length of 350bp. It is highly likely that the vast majority of authentic cCREs are just components of long CRMs, because even the longest cCREs (350bp) is shorter and the shortest VISTA enhancer (428bp). The highly uniform lengths of the predicted cCREs also indicate the limitation of the underlying prediction pipeline[26]. The distribution of GeneHancer enhancers oscillates with a period of 166bp (Figure 6E), which might be an artifact of the underlying algorithm for combining results from multiple sources [55]. Moreover, with a mean length of 1,488bp, GeneHancer enhancers are shorter than the VISTA enhancers (with a mean length 2,049bp) (Figure 6E). EnhancerAtlas enhancers also have a shorter mean length (680bp) than VISTA enhancers (3049bp) (Figure 6E). Our predicted CRMs at p-value <0.05 have a mean length of 1,162bp, thus also are shorter than that of VISTA enhancers (Figure 6E). However, as we indicated earlier, with a more stringent p-value cutoff 5×10^−6^, the resulting 428,628 predicted CRMs have almost an identical length distribution as the VISTA enhancers (Figure 3E). Taken together, these results unequivocally indicate that our predicted CRMs are much more accurate than the three state-of-the-art predicted enhancer/cCRE sets for both the nucleotide positions and lengths of CRMs/enhancers/cCREs.

### At least half of the human genome might code for CRMs

What is the proportion of the human genome coding for CRMs and TFBSs? The high accuracy of our predicted CRMs and constituent TFBSs might well position us to more accurately address this interesting and important, yet unanswered question[116, 117]. To this end, we took a semi-theoretic approach. Specifically, we calculated the expected number of true positives and false positives in the CRMCs in each non-overlapping *S*_*CRM*_ score interval based on the predicted number of CRMCs and the density of *S*_*CRM*_ scores of Null CRMCs in the interval (Figure 7A), yielding 1,383,152 (98.45%) expected true positives and 21,821 (1.55%) expected false positives in the CRMCs (Figure 7B). The vast majority of the 21,821 expected false positive CRMCs have a low *S*_*CRM*_ score < 4 (inset in Figure 7A) with a mean length of 28 bp, comprising 0.02% (21,821×28/3,088,269,832) of the genome and 0.05% (0.0002/0.4403) of the total length of the CRMCs, i.e., a FDR of 0.05% for nucleotide positions (Figure 7C). On the other hand, as the CRMCs miss 2.49% of ClinVar SNPs in the covered genome regions (Figure 5A), the FNR of partitioning the genome in CRMCs and non-CRMCs would be < 2.49%(1-0.40)=1.49%, given the proportion of neutrality of 0.4 for the unrecalled ClinVar SNPs (Figure 5B). False negative CRMCs would make up 0.67% of the genome and 1.99% of the total length of the non-CRMCs, meaning a false omission rate (FOR) of 1.99% for nucleotide positions (Figure 7C). Hence, true CRM positions in the covered regions would make up 44.68% (44.03%-0.02%+0.67%) of the genome (Figure 7C). In addition, as we argued earlier, the CRMC density in the uncovered 22.53% genome regions is about 80.04% of that in the covered regions, thus, CRMCs in the uncovered regions would be about 10.40% (0.2253 × 0.4468×0.8004/0.7747) of the genome (Figure 7C). Taken together, we estimated about 55.08% (44.68%+10.40%) of the genome to code for CRMs, for which we have predicted 79.90% [(44.03-0.02)/55.08]. Moreover, as we predicted that about 40% of CRCs are made up of TFBSs (Figure 3C), we estimated that about 22.03% of the genome might encode TFBSs. Furthermore, assuming a mean length 1,162bp for CRMs (the mean length of our predicted CRMs at p-value <0.05), and a mean length of 10bp for TFBSs (Figure 2D), we estimated that the human genome would encode about 1,463,872 CRMs (3,088,269,832×0.5508/1,162) and 67,034,584 TFBSs (3,088,269,832×0.2203/10).

**Figure 7.**
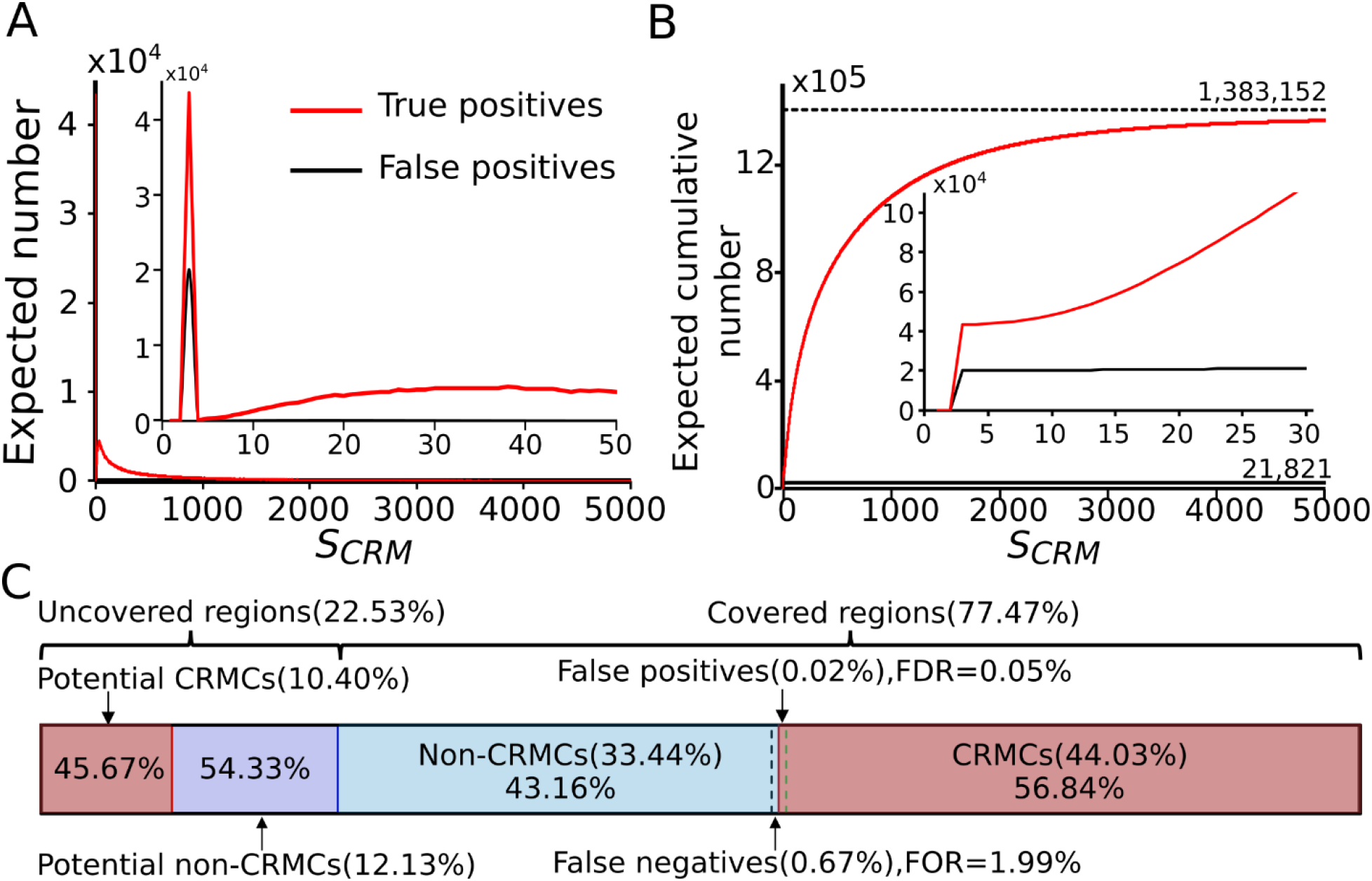
Estimation of the portion of the human genome encoding CRMs. A. Expected number of true positive and false positive CRMCs in the predicted CRMCs in each one-unit interval of the *S*_*CRM*_ score. The inset is a blow-up view of the axes defined region. B. Expected cumulative number of true positives and false positives with the increase in *S*_*CRM*_ score cutoff for predicting CRMs. The inset is a blow-up view of the axes defined region. C. Proportions of the genome that are covered and uncovered by the extended binding peaks and estimated proportions of CRMCs in the regions. Numbers in the braces are the estimated proportions of the genome being the indicated sequence types, and numbers in the boxes are proportions of the indicated sequence types in the covered regions or the uncovered regions, so they are summed to 1.

## Discussion

Identification of all functional elements, in particular, CRMs in genomes has been the central task in the postgenomic era, and enormous CRM function-related data have been produced to achieve the goal[23, 118]. Although great progresses have been made to predict CRMs in the genomes [26, 53, 55, 59, 119] using these data, most existing methods attempt to predict cell/tissue specific CRMs using CA and multiple tracks of histone marks collected in the same cell /tissue types[26, 48, 55, 59, 114]. These methods are limited by the scope of applications[26, 48, 114], low resolution of predicted CRMs[26, 59], lack of constituent TFBS information[26, 59], and high FDRs[68](Figure 6D). To overcome these limitations, we proposed a different approach to first predict a cell type agnostic or static map of CRMs and constituent TFBBs in the genome[46, 47] by identifying repeatedly cooccurring patterns of motifs found in appropriately expanded binding peaks in a large number of TF ChIP-seq datasets for different TFs in various cell/tissue types. Since it is mainly TFBSs in a CRM that define its structure and function, it not surprising that TF ChIP-seq data are a more accurate predictor of CRMs than CA and histone mark data[52, 68, 70]. Therefore, our approach might hold promise for more accurate predictions of CRMs and constituent TFBSs, notwithstanding computational challenges. Once a map of CRMs and constituent TFBBs in the gnome is available, functions of CRMs and constituent TFBSs in cell/tissue types could be studied in a more focused and cost-effective ways. Another advantage of our approach is that we do not need to exhaust all TFs and all cell/tissue types of the organism in order to predict most, if not all, of CRMs and constituent TFBBSs in the genome as we demonstrated earlier[46, 47], because CRMs are often repeatedly used in different cell/tissue types, developmental stages and physiological homeostasis[1]. Moreover, by appropriately extending the binding peaks in each dataset, we could largely increase the chance to identify cooperative motifs and full-length CRMs, thereby increasing the power of existing data, thereby further reducing the number of datasets needed as we have demonstrated in this and previous studies [46, 47]. We might only need a large but limited number of datasets to predict most, if not all, CRMs and TFBSs in the genome, as predicted UMs and CRMs enters a saturation phase when more than few hundreds of datasets were used for the predictions as we showed earlier [46]. Our earlier application of the approach resulted in very promising results in the fly[47] and human[46] genomes even using a relatively small number of strongly biased datasets available then. However, the earlier implementations were limited by computational inadequacies of underlying algorithms to find and integrate motifs in ever increasing number of large TF ChIP-seq datasets in mammalian cell/tissues[46, 47]. In this study, we circumvent the limitations by developing the new pipeline dePCRM2 based on an ultrafast and accurate motif finder ProSampler, an efficient motif pattern integration method, and a novel CRM scoring function that captures essential features of full-length CRMs.

Remarkably, dePCRM2 enables us to partition the 77.47% genome regions covered by the extended binding peaks in 6,079 TF ChIP-seq datasets into two exclusive sets, i.e., the CRMCs and non-CRMCs. Multiple pieces of evidence strongly suggest that the partition might be highly accurate. First, the vast majority of the CRMCs are unlikely predicted by chance as suggested by their small p-values (Figure 3B). Second, even the subset of the CRMCs with the lowest *S*_*CRM*_ scores ((0,1]) are under stronger evolutionary constraints than the non-CRMCs (Figures 4B and S4B), indicating that even these low-scoring CRMCs are more likely to be functional than non-CRMCs, not to mention CRMCs with higher *S*_*CRM*_ scores that are under stronger evolutionary constraints (Figures 4C, 4D, S4C and S4D). Third, the vast majority of the CRMCs are under similarly strong evolutionary constraints, and a subset of the CRMCs with high *S*_*CRM*_ scores evolve like the ultra-conserved, development-related VISTA enhancers with trimodal GERP score distributions (Figures 4A and S4A). Fourth, all experimentally validated VISTA enhancers and almost all (97.51%) of well-documented ClinVar SNPs in the covered genome regions are recalled by the CRMCs (Figure 5A), indicating that the CRMCs have a very low FNR. Finally, our simulation studies indicate that the CRMCs have a very low FDR of 0.05%, and the non-CRMCs have a low FOR of 1.99% (Figure 7C), strongly suggesting that both sensitivity and specificity of our predicted CRMs are very high. To the best of our knowledge, we are the first to accurately partition large regions (77.47**%)** of the genome into a set (CRMCs) that are highly likely to be *cis*-regulatory, and a set (non-CRMCs) that are highly unlikely to be *cis*-regulatory.

Accurate prediction of the length of CRMs is also critical, but it appeared to be a difficult problem as evidenced by the peculiar distributions of the lengths of GeneHancer enhancers and cCREs (Figure 6E). Although 44.26% (621,841) of our predicted 1,404,973 CRMCs are shorter than the shortest (428bp) VISTA enhancer, and thus are likely CRM components, they comprise only 7.42% of the total length of the CRMCs. The remaining 55.74% (783,132) of the CRMCs comprising 92.58% of the total length of the CRMCs, are longer than the shortest (428bp) VISTA enhancer, and thus are likely full-length CRMs. Therefore, the vast majority of the predicted CRMC positions in the genome might be covered by full-length CRMs. Very short CRMCs tend to have small *S*_CRM_ scores and be under weak evolutionary constraints, and thus can be effectively filtered out using more stringent *S*_CRM_ cutoffs (Figures 3E, 4C and S4C). It has been shown that an enhancer’s length and evolutionary behavior are determined by its regulatory tasks [91], and conserved enhancers are active in development [98, 99], while fragile enhancers are associated with evolutionary adaptation [98]. CRMCs with different S_CRM_ cutoffs might belong to different functional types as indicated by their different evolutionary behaviors (4A, 4C, S4A and S4C) and length distributions (Figures 3E). For example, like VISTA enhancers, CRMs predicted at high S_CRM_ cutoffs tend to be longer (Figure 3E) and under stronger evolutionary constrains (Figures 4C and S4C), thus might be mainly involved in development, whereas CRMs predicted at lower S_CRM_ cutoffs tend to be shorter (Figure 3E) and under weaker evolutionary constrains (Figures 4C and S4C), thus might be mainly involved in non-development related functions. On the other hand, the failure to predict full-length CRMs of short CRM components might be due to insufficient data coverage on the relevant loci in the genome. This is reminiscent of our earlier predicted, even shorter CRMCs (mean length = 182bp) using a much smaller number and less diverse 670 datasets[46]. As we argued earlier[46] and confirmed here by the much longer CRMCs (mean length = 981bp) predicted using the much larger and more diverse datasets albeit still strongly biased to a few TFs and cell/tissue types (Supplementary Note). We anticipate that full-length CRMs of these short CRM components can be predicted using even larger and more diverse TF ChIP-seq data. Thus, efforts should be made in the future to increase the genome coverage and reduce data biases by including more untested TFs and untested cell types in the TF ChIP-seq data generation.

Interestingly, our predicted CRMs (at p-value < 0.05) achieve perfect (100.00%) and very high (97.43) sensitivity for recalling VISTA enhancers [79] and ClinVAR SNPs [103], respectively, but varying intermediate sensitivity ranging from 64.50% (for GWAS SNPs) to 88.77% (for FPs) for recalling other CRM function-related elements in the remaining eight datasets (Figure 5A). It appears that such varying sensitivity is due to varying FDRs ranging from 0% (for VISTA enhancers) to 35.5% (for GWAS SNPs) of the methods used to characterize the elements (Figure 5B). Our finding that DHSs, TASs, and histone mark (H3K4m1, H3K4m3 and H3K27ac) peaks have high FDRs for predicting CRMs is consistent with an earlier study showing that histone marks or CA were less accurate predictor of active enhancers than TF binding data[68]. Thus, it is not surprising that our predicted CRMs substantially outperforms the three state-of-the-art sets of predicted enhancers/cCREs, i.e., GeneHancer 4.14 [55], EnhancerAtals2.0 [59] and cCREs[26], both for recalling VISTA enhancers (we excluded GeneHancer enhancers for this evaluation since VISTA enhancers were a part of it) and ClinVar SNPs (Figure 6B) and for predicting the lengths of CRMs (Figure 6E), because these three sets were mainly predicted based on overlaps between multiple tracks of CA and histone marks in various cell/tissue type. Although great efforts have been made to improve the accuracy of EnhancerAtlas 2.0 enhancers[59], GeneHancer 4.14 enhancers[55] and cCREs[26], they still suffer quite high FDRs (49.14%, 29.28% and 23.12%, respectively).

Although dePCRM2 can predict functional states of CRMCs in a cell/tissue type that have original binding peaks overlapping the CRMs, it cannot predict the functional states of CRMs in the extended parts of the original binding peaks in a cell/tissue if the CRMs do not overlap any available binding peaks of all TFs tested in the cell/tissue type. However, the functional state of each CRM in the map in any cell/tissue type could be predicted based on overlap between the CRM and a single or few epigenetic mark datasets such as CA, H3K27ac and/or H3K4m3 data collected from the very cell/tissue type. Anchored by correctly predicted CRMs, these epigenetic marks could accurately predict the functional states of the CRMs[68]. Thus, our approach might be more cost-effective for predicting both a static map of CRMs as well as constituent TFBSs in the genome and their functional states in various cell/tissue types.

Remarkably, although originally called binding peaks is the strongly biased to few cell types and TFs (Supplementary Note), and the 6,092 TF ChIP-seq datasets cover only 40.96% of the genome, after moderately extending the binding peaks, we increased the genome coverage to 77.47%, an 89.14 % increase. Nucleotide positions of the extended parts of the peaks contribute 42.12% positions of the predicted CRMCs. Therefore, appropriate extension of called binding peaks in the datasets can substantially increase the power of available data. On the other hand, we abandoned 38.60% of positions covered by the original binding peaks, which might be nonfunctional as they evolve like the non-CRMCs (Figures 4A and S4A). Therefore, originally called binding peaks cannot be equivalent to CRMs or parts of CRMs as has also been shown earlier[87-89], and integration of multiple TF ChIP-seq datasets as demonstrated in this study is necessary for accurate genome-wide predictions of CRMs.

The proportion of the human genome that is functional is a topic under hot debate [109, 120] and a wide range from 5% to 80% of the genome has been suggested to be functional based on difference sources of evidence [23, 61, 116, 120, 121]. The major disagreement is for the proportion of functional NCSs in the genome, mainly CRMs, which have been coarsely estimated to comprise from 8% to 40% of the genome [109, 120]. Moreover, a wide range of CRM numbers from 400,000 [109] to more than a few million [23, 59] has been suggested to be encoded in the human genome. However, to our best knowledge, no estimate has been made on substantial evidence. Our predicted CRMCs cover 44.03% of the genome, which is lower than EnhancerAtlas enhancers (58.99%)[59] do. The much higher accuracy of our predicted CRMs suggests that cCREs (7.9%)[26] and GeneHancer enhancer might underpredict, whereas EnhancerAtlas 2.0 might overpredict CRMs. Based on the estimated FDR and FNR of the predicted CRMCs and non-CRMCs as well as the estimated density of CRMs in the uncovered regions relative to the covered regions (Figure 7C), we estimated that about 55.08% and 22.03% of the genome might code for CRMs and TFBSs, respectively, which encode about 1.46 million CRMs and 67 million TFBSs. Therefore, the number of our predicted CRMs is almost four times more than an earlier estimate of 400,000 [109], and they are encoded by a higher (55.08%) proportion of the genome than earlier thought 40%[109, 120]. We estimated that our true positive CRMs cover 44.01% (44.03-0.02) of the genome, therefore, we might have predicted 79.90 % (44.01/55.08) CRM positions encoded in the genome. In summary, it appears that the *cis*-regulatory genome is more prevalent than originally thought.

## Conclusions

We have developed a new highly accurate and scalable pipeline dePCRM2 for predicting CRMs and constituent TFBSs in large genomes by integrating a large number of TF ChIP-seq datasets for various TFs in a variety of cell/tissue types of the organisms. Applying dePCRM2 to all available ∼6,000 TF ChIP-seq datasets, we predicted an unprecedentedly complete, high resolution map of CRMs and constituent TFBSs in 77.47% of the human genome covered by extended binding peaks of the datasets. Evolutionary and experimental data suggest that dePCRM2 achieves very high prediction sensitivity and specificity. Based on the predictions, we estimated that about 55% and 22% of the genome might code for CRMs and TFBSs, encoding about 1.46 million CRMs and 67 million TFBSs, respectively; for both of which we predicted about 80%. Therefore, the *cis*-regulatory genome is more prevalent than originally thought. With the availability in the future of more diverse and balanced data covering more regions of the genome, it is possible to predict a more complete map of CRMs and constituent TFBSs in the genome.

## Materials and Methods

### Datasets

We downloaded 6,092 TF ChIP-seq datasets from the Cistrome database[29]. The binding peaks in each dataset were called using a pipeline for uniform processing[29]. We filtered out binding peaks with a read depth score less than 20. For each binding peak in each dataset, we extracted a 1,000 bp genome sequence centering on the middle of the summit of the binding peak. We downloaded 976 experimentally verified enhancers and 942 negatively validated regions (NVRs) from the VISTA Enhancer database[79], 424,622 ClinVar SNPs from the ClinVar database[103], 32,689 enhancers[101] and 184,424 promoters[100] from the FANTOM5 project website, 91,369 GWAS SNPs from GWAS Catalog[102], and 122,468,173 DHSs in 1,353 datasets (Table S4), 29,520,736 transposase-accessible sites (TASs) in 1,059 datasets (Table S5), 99,974,447 H3K27ac peaks in 2,539 datasets (Table S6), 77,500,232 H3K4me1 peaks in 1,210 datasets (Table S7), and 70,591,888 H3K4me3 peaks in 2,317 datasets (Table S8) from the Cistrome database[29].

### Measurement of the overlap between two different datasets

To evaluate the extent to which the binding peaks in two datasets overlap with each other, we calculate an overlap score *S*_0_ (*d*_*i*_, *d*_*j*_) between each pair of datasets *d*_*i*_ and *d*_*j*_, defined as,

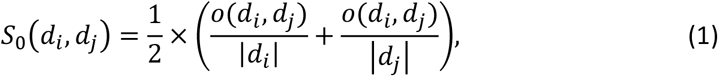

where *o*(*d*_*i*_, *d*_*j*_) is the number of binding peaks in *d*_*i*_ and *d*_*j*_ that overlap each other by at least one bp.

### Parameters for accuracy evaluation

We use the following definitions to evaluate the accuracy of datasets and predictions. *S*ensitivity = recall 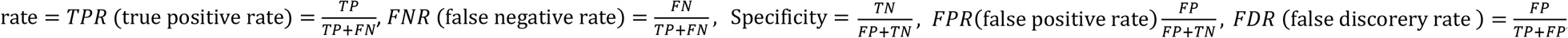, and 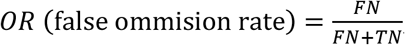, where TP is true positives; FN, false negatives; FP, false positives; and TN, true negatives.

### The dePCRM2 pipeline

**Step 1:** Find motifs in each dataset using ProSampler[72](Figures 1A and 1B).

**Step 2**. Compute pairwise motif co-occurring scores and find co-occurring motif pairs (CPs): As True motifs are more likely to co-occur in the same sequence than spurious ones, to filter out false positive motifs, we find overrepresented CPs in each dataset (Figure 1C). Specifically, for each pair of motifs *M*_*d*_(*i*) and *M*_*d*_(*j*) in each data set *d*, we compute their co-occurring scores *S*_*c*_ defined as,

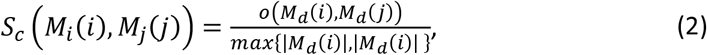

where |*M*_*d*_ (*i*)| and |*M*_*d*_ (*j*)| are the number of binding peaks containing TFBSs of motifs *M*_*d*_ (*i*) and *M*_*d*_(*j*), respectively; and *o*(*M*_*d*_(*i*), *M*_*d*_(*j*)) the number of binding peaks containing TFBSs of both the motifs in *d*. We identify CPs with an *S*_*c*_ ≥ *β*. We choose *β* such that the component with the highest scores in the trimodal distribution *S*_*c*_ is kept (Figures 1C and 2B) (by default *β* = 0.7).

**Step 3**. Construct a motif similarity graph and find unique motifs (UMs): We combine highly similar motifs in the CPs from different datasets to form a UM presumably recognized by a TF or highly similar TFs of the same family/superfamily[122]. Specifically, for each pair of motifs *M*_*a*_(*i*) and *M*_*b*_ (*j*) from different datasets *a* and *b*, respectively, we compute their similarity score *S*_*s*_ using our SPIC[123] metric. We then build a motif similarity graph using motifs in the CPs as nodes and connecting two motifs with their *S*_*s*_ being the weight on the edge, if and only if (iff) *S*_*s*_ >*β* (by default, *β =*0.8, Figure 1D). We apply the Markov cluster (MCL) algorithm [124] to the graph to identify dense subgraphs as clusters. For each cluster, we merge overlapping sequences, extend each sequence to a length of 30bp by padding the same number of nucleotides from the genome to the two ends, and then realign the sequences to form a UM using ProSampler[72](Figure 1D).

**Step 4**. Construct the interaction networks of the UMs/TFs: TFs tend to repetitively cooperate with each other to regulate genes in different contexts by binding to their cognate TFBSs in CRMs. The relative distances between TFBSs in a CRM often do not matter (billboard model), but sometimes they are constrained by the interactions between cognate TFs (enhanceosome model) [74-76]. To model essential features of both scenarios, we compute an interaction score between each pair of UMs, *U*_*i*_ and *U*_*j*_, defined as,

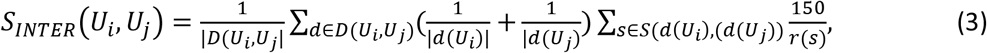

where *D*(*U*_*i*_, *U*_*j*_) is the datasets in which TFBSs of both *U*_*i*_ and *U*_*j*_ occur, *d*(*U*_*k*_) the subset of dataset *d*, containing at least one TFBS of *U*_*k*_, *S*(*d*(*U*_*i*_), (*d*(*U*_*j*_)) the subset of *d* containing TFBSs of both *U*_*i*_ and *U*_*j*_, *and r*(*s*) the shortest distance between any TFBS of *U*_*i*_ and any TFBS of *U*_*j*_ in a sequence *s*. We construct UM/TF interaction networks using the UMs as nodes and connecting two nodes with their *S*_INTER_ being the weight on the edge (Figure 1E). Therefore, the S_INTER_ score allows flexible adjacency and orientation of TFBSs in a CRM (billboard model) and at the same time, it rewards motifs with binding sites co-occurring frequently in a shorter distance in a CRM (enhanceosome model), particularly within a nucleosome with a length of about 150bp[74, 75, 125].

**Step 5**. Partition the covered genome regions into a CRM candidate (CRMC) set and a non-CRMC set: We project TFBSs of each UM back to the genome, and link two adjacent TFBSs if their distance *d* ≤ 300bp (roughly the length of two nucleosomes). The resulting linked DNA segments are CRMCs, while DNA segments in the covered regions that cannot be linked are non-CRMCs (Figure 1F).

**Step 6**. Evaluate each CRMC: We compute a CRM score for a CRMC containing *n* TFBSs (*b*_1_, *b*_2_, *…, b*_n_), defined as,

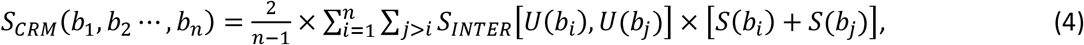

where *U*(*b*_*k*_) is the UM of TFBS *b*_*k*_, *S*_*INTER*_[*U*(*b*_*i*_), *U*(*b*_*j*_)] the weight on the edge between *U*(*b*_*i*_) and *U*(*b*_*j*_), in the interaction networks, and *S*(*b*_*k*_) the score of *b*_*k*_ based on the position weight matrix (PWM) of *U*(*b*_*k*_). Only TFBSs with a positive score are considered. Thus, *S*_*CRM*_ considers the number of TFBSs in a CRMC, as well as their quality and strength of all pairwise interactions.

**Step 7**. Predict CRMs: We create the Null interaction networks by randomly reconnecting the nodes with the edges in the interaction networks constructed in Step 4. For each CRMC, we generate a Null CRMC that has the same length and nucleotide compositions as the CRMC using a third order Markov chain model[72]. We compute a S_CRM_ score for each Null CRMC using the Null interaction networks, and the binding site positions and PWMs of the UMs in the corresponding CRMC. Based on the distribution of the S_CRM_ scores of the Null CRMCs, we compute an empirical p-value for each CRMC, and predict those with a p-value smaller than a preset cutoff as CRMs in the genome (Figure 1G).

**Step 8**. Prediction of the functional states of CRMs in a given cell type: For each predicted CRM at p-value <0.05, we predict it to be active in a cell/tissue type, if its constituent binding sites of the UMs whose cognate TFs were tested in the cell/tissue type overlap original binding peaks of the TFs; otherwise, we predict the CRM to be inactive in the cell/tissue type. If the CRM does not overlap any binding peaks of the TFs tested in the cell/tissue type, we assign its functional state in the cell/tissue type “TBD” (to be determined).

### Generation of control sequences for validation

To create a set of matched control sequences for validating the predicted CRMs using experimentally determined elements used in Figure 5A, for each predicted CRMC, we produced a control sequence by randomly selecting a sequence segment with the same length as the CRMC from the genome regions covered by the extended binding peaks. To calculate the S_CRM_ score of a control sequence, we assigned it the TFBS positions and their UMs according to those in the counterpart CRMC. Thus, the control set contains the same number and length of sequences as in the CRMCs, but with arbitrarily assigned TFBSs and UMs.

## Authors’ contributions

ZS conceived the project. ZS and PN developed the algorithms and PN carried out all computational experiments and analysis. ZS and PN wrote the manuscripts. All authors read and approved the final manuscript.

## Ethics approval and consent to participate

Not applicable.

