## Supplemental Figures for "Accurate prediction of *cis*-regulatory modules reveals a prevalent regulatory genome of humans"

Supplementary Figure S1

A

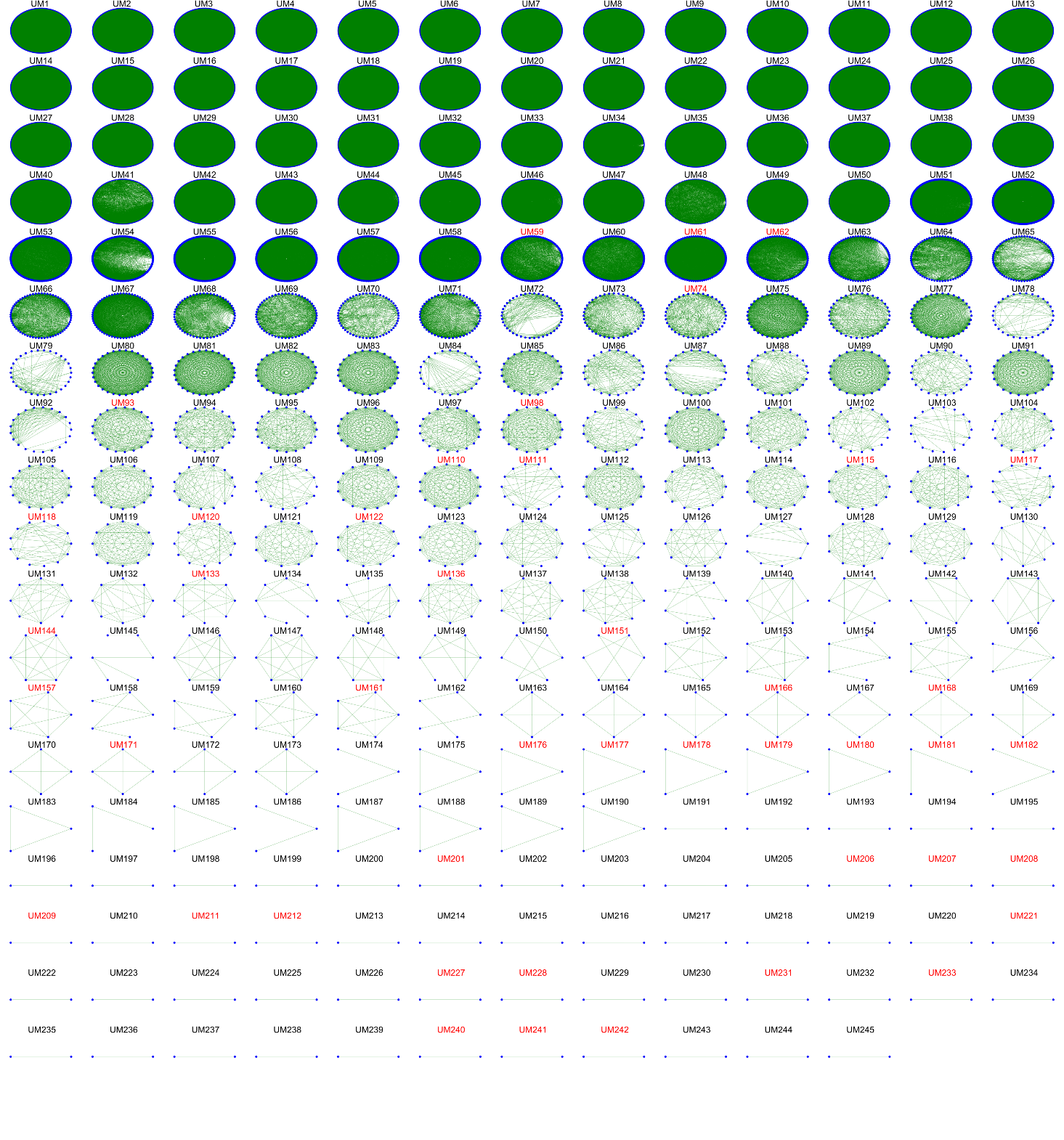

(B)

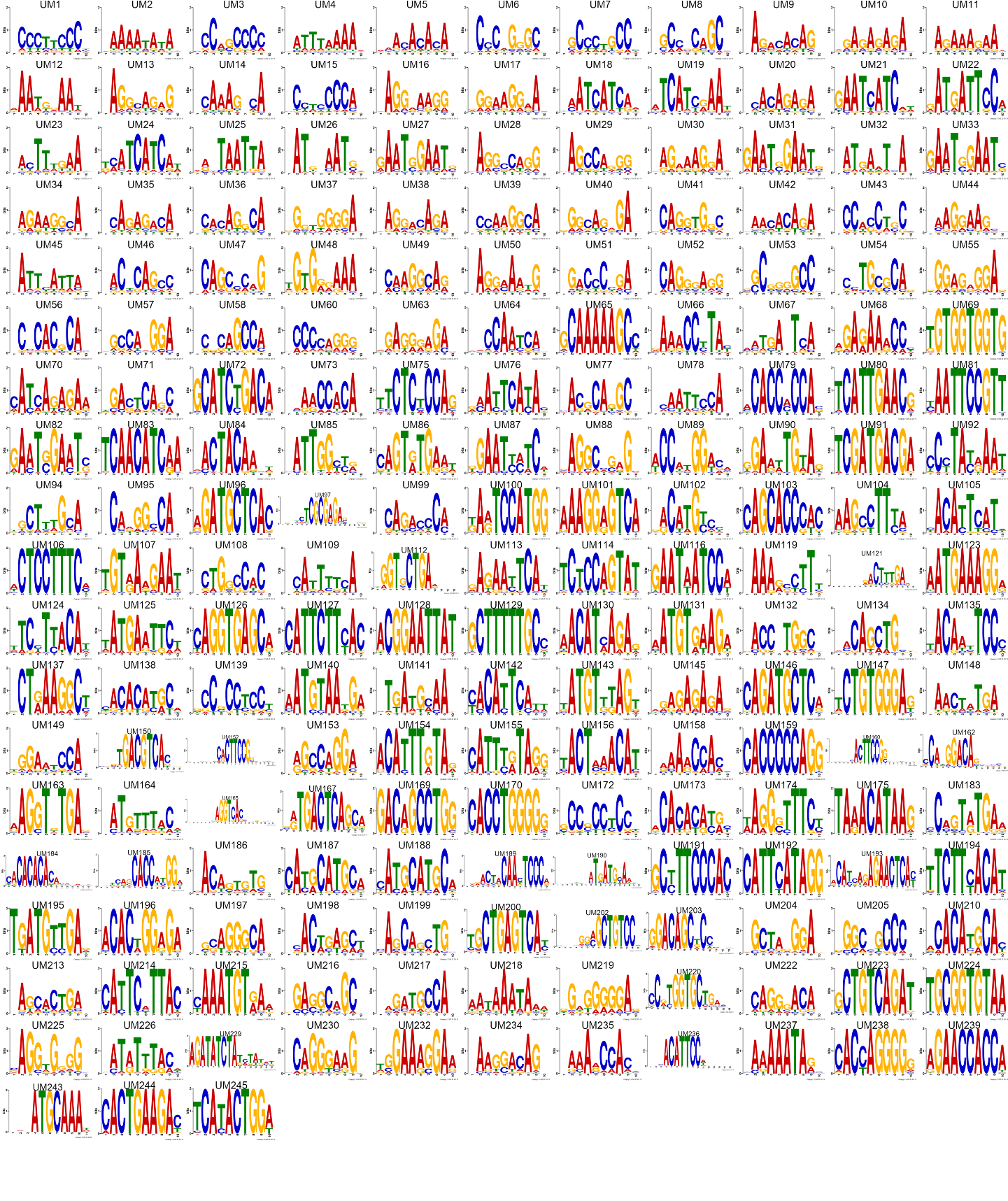

Figure S1. Prediction of UMs. **A.** Similarity graphs of member motifs in the 245 motif clusters. In each graph, a node in blue represents a member motif of the cluster, and two member motifs are connected by an edge in green if their similarity is greater than 0.8 (SPIC score). Clusters with the names in RED font are those in which a UM cannot be found. **B.** Logos of the 201 UMs found in the corresponding clusters.

Supplementary Figure S2

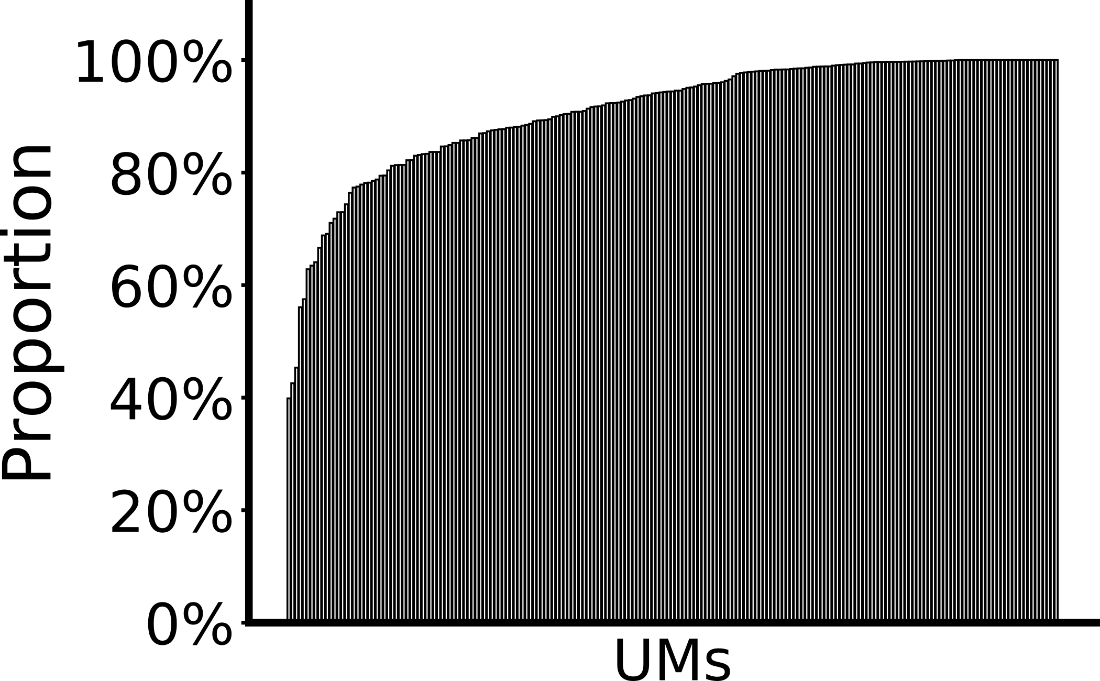

Figure S2. Proportion of sequences of the member motifs of a UM in which binding sites were found. UMs are sorted in ascending order of the proportion.

Supplementary Figure S3

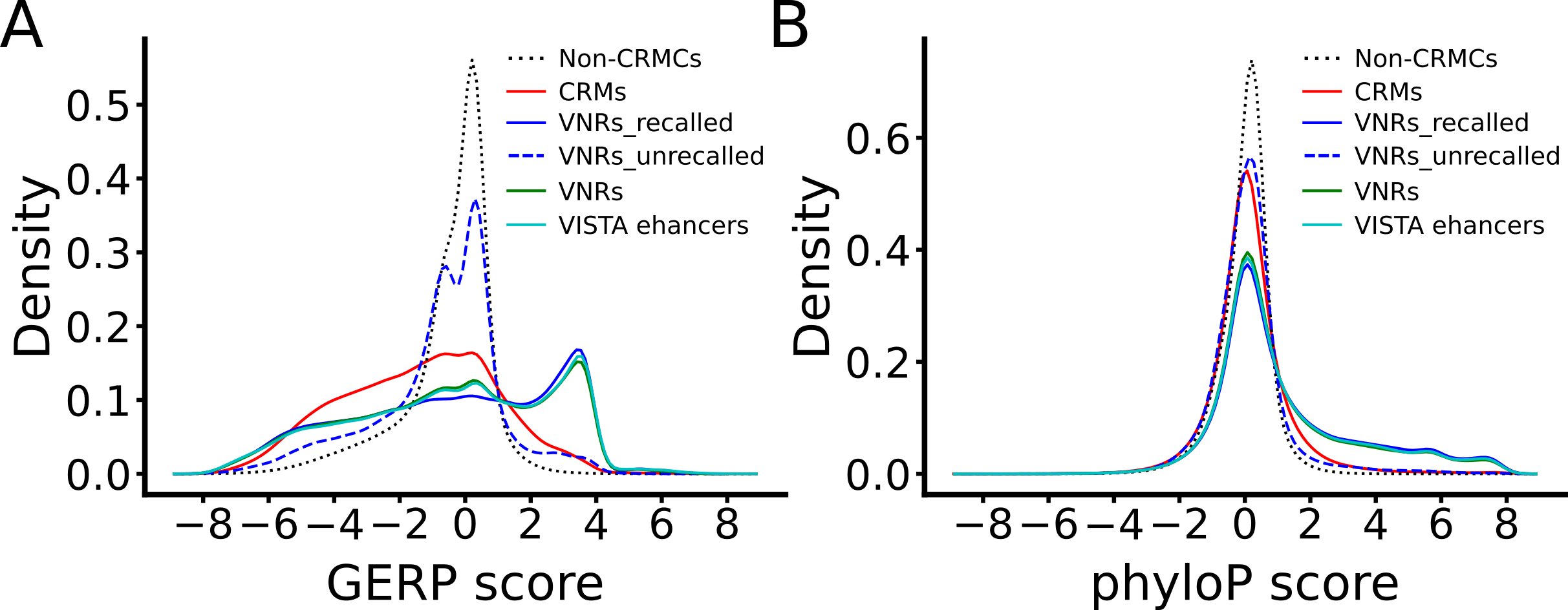

Figure S3. Comparison of evolutionary behaviors of VISTA enhancers and VNRs. **A.** Distributions of the GERP scores of nucleotide positions of the predicted CRMCs, non-CRMCs, VISTA enhancers, VNRs, recalled NVRs and unrecalled NVRs. **B.** Distributions of the phyloP scores of nucleotide positions of the predicted CRMCs, non-CRMCs, VISTA enhancers, VNRs, recalled NVRs and unrecalled NVRs.

Supplementary Figure S4

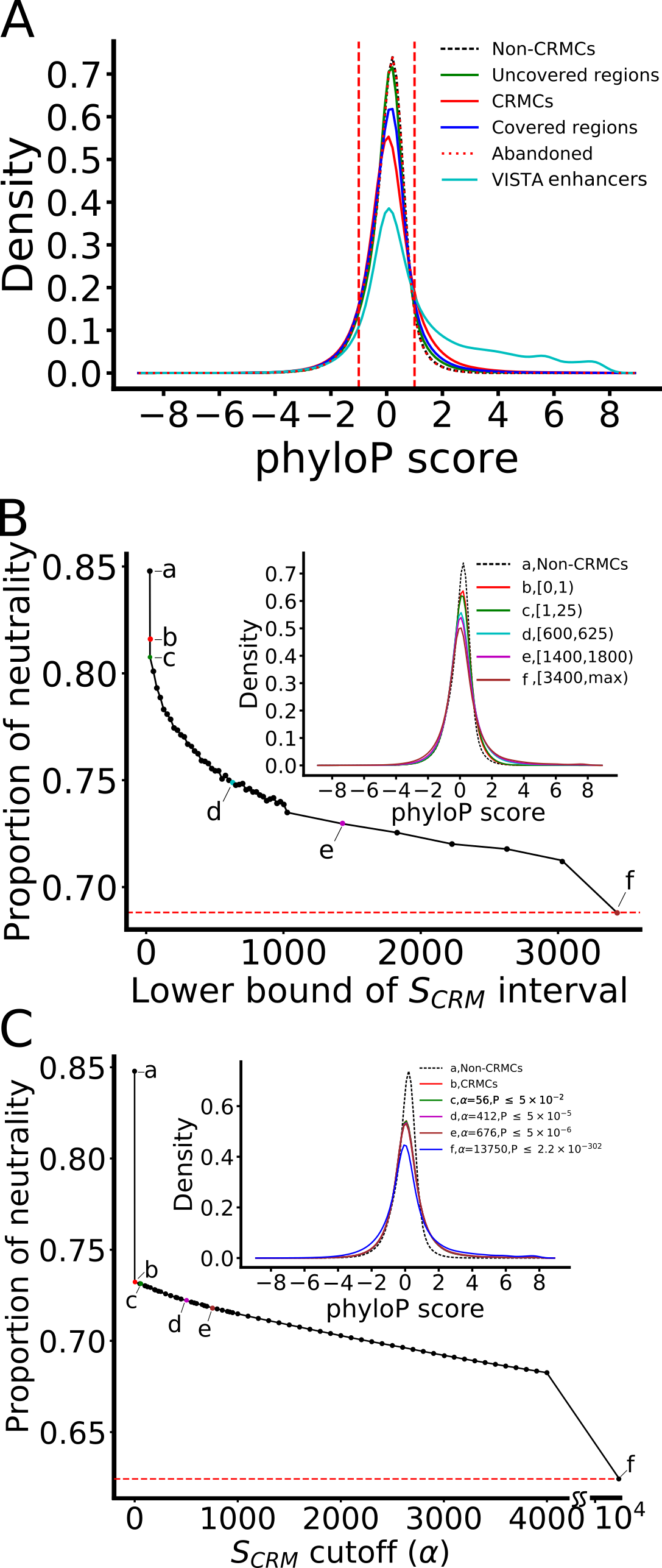

Figure S4. CRMCs and non-CRMCs in NESs show different evolutionary behaviors measured by phyloP scores. A. Distributions of the phyloP scores of nucleotide positions of VISTA enhancers, CRMCs, non-CRMCs, abandoned genome regions covered by original binding peaks, genome regions covered by extended binding peaks and genome regions uncovered by extended binding peaks.  The area under the density curves in the score interval [-1, 1] is defined as the proportion of neutrality of the sequences. B. Proportion of neutrality of CRMCs with a S_CRM_ score in different intervals in comparison with that of the non-CRMCs (a). The inset shows the distributions of the phyloP scores of the non-CRMCs and CRMCs with S_CRM_ scores in the intervals indicted by color and letters.  C. Proportion of neutrality of CRMs predicted using different S_CRM_ score cutoffs and associated p-values in comparison with those of the non-CRMCs (a) and CRMCs (b).  The inset shows the distributions of the phyloP scores of the non-CRMCs, CRMCs and the predicted CRMs using the S_CRM_ score cutoffs and p-values indicated by color and letters. The dashed lines in B and C indicate the saturation levels.

Supplementary Figure S5

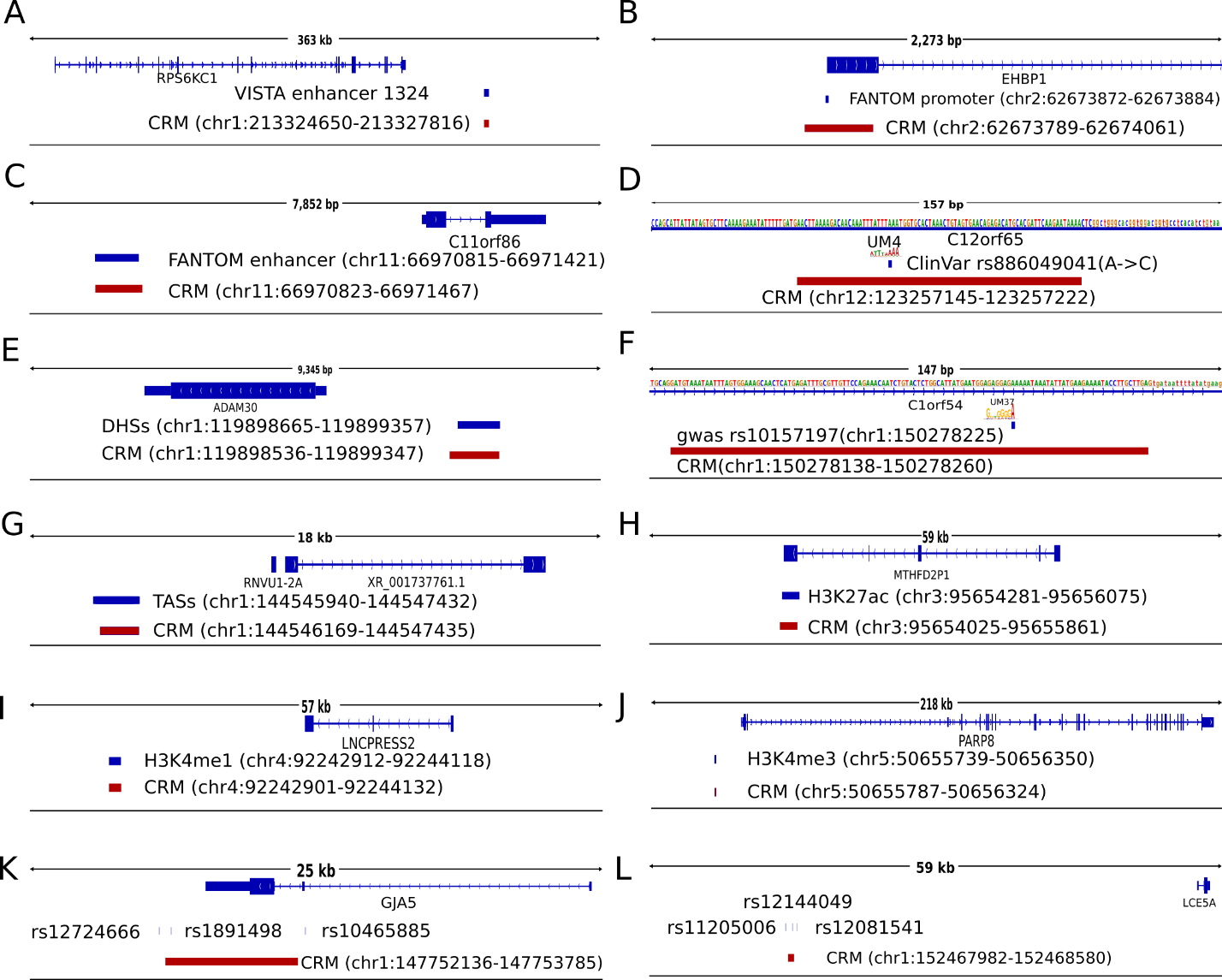

Figure S5. Examples of predicted CRMs that recover experimentally determined sequence elements. **A**. A CRM (chr1:213324650-2733277816) recovers VISTA enhancer 1324 downstream of gene *RPS6K1*. **B.** A CRM (chr2:62673789-62674061) recovers a FANTOM5 promoter (chr2: 62673872-62673884) for gene EHBP1. **C.** A CRM (chr11:66970823-66971467) recovers a FANTOM5 enhancer (chr11: 66970815-66971421) upstream of gene *C11orf86*. **D.** A CRM (chr12:123257145-123257222) recovers a ClinVar point mutant rs886049041(A>C) in an intron of gene *C12orf65*, related to combined oxidative phosphorylation deficiency. The SNP is located in a critical position of UM4. **E.** A CRM (chr1:119898536-119899347) recovers a DHS (chr1: 119898665-119899357) upstream of gene *ADAM30.* **F.** A CRM (chr1:150278138-150278260) recovers a GWAS SNP rs10157197 (chr1:150278225) in an intron of gene *C1orf54*. The SNP is located in a critical position of UM37. **G.** A CRM (chr1:114546169-144547435) recovers a TAS (chr1:114545940-144547432) is located upstream of gene RNVU1-2A. **H.** A CRM (chr3:95654025-95655861) recovers a H3K27ac peak located downstream of the gene MTHFD2P1. **I.** A CRM (chr4:92242901-92244132) recovers a H3K4me1 peak (chr4:92242912-92244118) downstream of the gene LNCPRESS2. **J.** A CRM (chr5:50655787-50656324) recovers a H3K4me3 peak (chr5:50655739-50656350) upstream of the gene PARP8. **K.**A CRM (chr1:147752136-147753785) recovers GWAS SNP rs1891498 upstream of gene *GJA5*, while two unrecovered GWAS SNPS rs12724666 and rs10465885 located upstream of and in an intron of the gene, respectively, are in LD with rs1891498. **L.** A CRM (chr1:152467982-152468580) recovers a GWAS SNP rs12144049 upstream of gene *LCE5A,* while two unrecovered GWAS SNPS rs11205006 and rs12081541 located upstream and downstream of rs12144049, respectively, are in LD with it.
