## Supplemental Notes for "Accurate prediction of *cis*-regulatory modules reveals a prevalent regulatory genome of humans"

**Supplementary Note**

**Extended binding peaks in different datasets have extensive overlaps**

After filtering out low-quality peaks in the 6,092 ChIP-seq datasets (Table S1), we ended up with 6,070 non-empty datasets for 779 TFs in 2,631 cell/tissue types. The datasets are strongly biased to few cell types (Figure N1A). For example, 532, 475 and 309 datasets were collected from mammary gland epithelium, colon epithelium, and bone marrow erythroblast, respectively, while only one dataset was generated from 129 cell/tissue types, including heart embryonic fibroblast, fetal skin fibroblast, and bone marrow haematopoietic progenitor, and so on (Figure N1A and Table S1). The datasets also are strongly biased to few TFs (Figure N1B and Table S1). For example, 370 and 263 datasets were collected for TFs CTCF and ESR1, respectively, while just one dataset was produced for 324 TFs, such as MSX2, RAX2, and MYNN, and so on. The number of called binding peaks in a dataset is highly varying, ranging from two to 100,539, with a mean of 19,314 (Figure N1C). For instance, datasets for STAT1 and NR3C1 have the smallest number of two binding peaks in HeLa-S3 and HEK293 cells, respectively, while datasets for CEBPB, BRD4 and FOXA2 have the largest number of 115,776, 99,646, and 99,512 binding peaks in HepG2, U87, and Mesenchymal Stem Cells, respectively. The highly varying numbers of binding peaks in the datasets suggest that different TFs might bind a highly varying number of sites in the genomes of different cell/tissue types. However, some datasets with very few binding peaks might be resulted from technical artifacts, thus are of low quality (see below), even though they passed our first quality filter. The lengths of binding peaks in the datasets range from 75 to 10,143bp, with a mean of 300bp (Figure N1D), and 99.12% of binding peaks are shorter than 1,000pb. All the binding peaks in the 6,070 datasets cover a total of 1,265,474,520bp (40.98%) of the genome (3,088,269,832bp). For each binding peak in each dataset, we extracted a 1,000bp genome sequence centering on the middle of its binding summit, thereby extending the lengths for most binding peaks. We have shown that extension of binding peaks to 500~1,000bp could substantially increase the chance of finding TFBSs of cooperators of the ChIP-ed TFs, while the introduced noise had a little effect on identifying the primary motifs of ChIP-ed TFs[1]. The extended binding peaks contain a total of 115,710,048,000bp, which is 37.5 times the size of the genome, but cover only 2,392,488,699bp (77.47%) of the genome, leaving the remaining 22.53% of the genome uncovered, indicating that they have extensive overlaps. Remarkably, by extending the originally called binding peaks, we increased the coverage of genome by 89.04% (77.47% vs 40.98%). However, we may not know the functional states of some predicted binding sites in extended parts of the original binding peaks from a cell/tissue type if no binding peaks for other TFs tested in the cell/tissue type overlap the extended parts. We trade this drawback for a more complete prediction of the CRMs and TFBSs in the genome[2]. As stated earlier, we predict CRMs and constituent TFBSs based on overlapping patterns between datasets for ChIP-ed cooperative TFs, thus, we evaluated the extent to which the extended binding peaks in different datasets overlap one another. To this end, we hierarchically clustered the 6,070 datasets using an overlap score between each pair of the datasets (formula 1). As shown in Figure N1E, there are extensive distinct overlapping patterns among the datasets. As expected, clusters are formed by datasets for largely the same TF in different cell/tissue types, and/or by datasets for different TFs that are known or potential collaborators in transcriptional regulation. For instance, a cluster is formed by 1, 1 and 48 datasets for RAD21, SMC3 and CTCF, respectively, in certain cell/tissue types (Figure N1F). It is well-known that RAD21, SMC3 and CTCF are core subunits of the cohensin complex, and are widely colocalized in mammalian genomes[3]. In another example (Figure N1G), a cluster is formed by 50 datasets for 45 TFs in various cell/tissue types. Multiple sources of evidence indicate that these 45 TFs have extensive physical interactions for DNA binding and transcriptional regulation in certain cell/tissue types (Figure N1H)[4, 5]. For example, it has been shown that LEF1 interacts with the TGF beta activating regulator SMAD4 [6] , E2F4 helps to recruit PML to the TBX2 promoter [7], and FOXK 1 and TP53 can form a distinct protein complex on and off chromatin[8]. These overlapping patterns between the extended binding peaks in the datasets warrant us to predict CRMs and constituent TFBSs in the covered genome regions[2, 9]. In addition, we collected 10 sets of experimentally determined CRM function-related elements (Materials and Methods) and found that they are enriched in the covered 77.47% genome regions relative to the uncovered 22.53% regions, including 785 (80.43%) VISTA enhancers[10], 402,730 (94.84%) of ClinVar SNPs[11], 181,436 (98.38%) FANTOM5 promoters (FPs)[12], 32,029 (97.98%) FANTOM5 enhancers (FEs)[13], 82,378 (90.16%) of GWAS SNPs[14], 121,075,184 (98.86%) DHSs[15] (Table S4), 29,195,778 (98.90%) TASs[15](Table S5), 98,297,240(98.32%) H3K27ac peaks[15](Table S6), 75,467,050 (97.38%) H3K4me1 peaks[15](Table S7), and 69,282,044(98.14%) H3K4me3 peaks[15](Table S8). These results indicate that the covered genome regions were better studied than the uncovered regions. We will evaluate the sensitivity of dePCRM2 to recall these elements at different $S_{CRM}$ scores and associated p-value cutoffs.

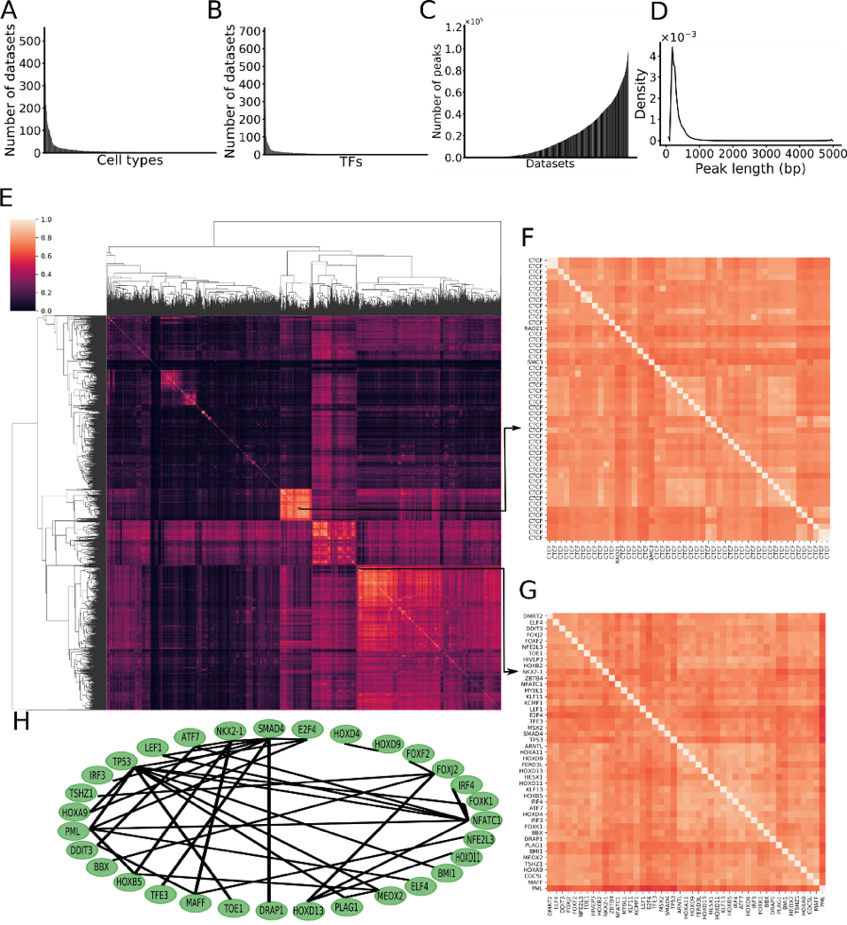

Figure N1. Properties of the datasets. A. Number of datasets collected in each cell/tissue types sorted in descending order. B. Number of datasets collected for each TF sorted in descending order. C. Number of peaks in each dataset sorted in ascending order. D. Distribution of the lengths of binding peaks in the entire datasets. E. Heatmap of overlaps of extended binding peaks between each pair of the datasets. F. A blowup view of the indicated cluster in E, formed by 48 datasets for CTCF in different cell/tissue types, as well as one dataset for each of its two collaborators, RAID21 and SMC3. G. A blowup view of the indicated cluster in E, formed by 50 datasets for 45 TFs. H. Known physical interactions between the 45 TFs whose 50 datasets form the cluster in E.
